## supporting information for "The Meta-Position of Phe^4^ in Leu-enkephalin Regulates Potency, Selectivity, Functional Activity, and Signaling Bias at the Delta and Mu Opioid Receptors"

#### Table of Contents

Table S1……………………………………………………………………….………… SI-2

Table S2………………………………………………………………………………..SI-3

Characterization of Peptides ........................................................................................SI-4

NMR Spectra of Peptides..............................................................................................SI-9

HPLC Chromatograms of Peptides..............................................................................SI-24

**Table S1: Efficacy of cAMP inhibiton of Meta-substituted Phe$^4$ Analogs of Leu-enkephalin at δOR and μOR.**

| **cAMP** | **δOR** | **μOR** |
|---|---|---|
| Compound | Efficacy (%+SD) | Efficacy (%+SD) |
| **1a** (F) | 95 ± 6 | 98 ± 6 |
| **1b** (Cl) | 94 ± 5 | 99 ± 7 |
| **1c** (Br) | 96 ± 4 | 104 ± 11 |
| **1d** (I) | 97 ± 4 | 94 ± 11 |
| **1e** (Me) | 92 ± 8 | 85 ± 10 |
| **1f** (OMe) | 98 ± 10 | 87 ± 5 |
| **1g** ($CF_3$) | 92 ± 8 | 90 ± 4 |
| **1h** (CN) | 100 ± 3 | 92 ± 13 |
| **1i** ($NO_2$) | 99 ± 6 | 105 ± 12 |
| **1j** (2-pyr) | 100 ± 8 | 99 ± 5 |
| **1k** (3-pyr) | 97 ± 8 | 100 ± 7 |
| **1l** (4-pyr) | 99 ± 10 | 99 ± 6 |
| DADLE | 97 ± 4 | 98 ± 6 |
| Leu-enkephalin | 100 | 100 ± 9 |
| DAMGO | 98 ± 4 | 100 |

**Table S2: LogR values for Meta-substituted Phe[4] Analogs of Leu-enkephalin at δOR and μOR in the cAMP and β-arrestin 2 (β-arr2) assays. 95% confidence intervals are presented between parentheses.**

| | **LogR (δOR)** | | **LogR (μOR)** | |
|---|---|---|---|---|
| Compound | cAMP | β-arr2 | cAMP | β-arr2 |
| **1a** (F) | 9.4<br>(9.1-9.6) | 8.1<br>(7.9-8.3) | 8.0<br>(7.7-8.3) | 5.8<br>(5.6-6.1) |
| **1b** (Cl) | 10.6<br>(10.3-10.9) | 9.0<br>(8.8-9.3) | 7.5<br>(7.1-7.8) | 7.2<br>(6.9-7.4) |
| **1c** (Br) | 10.5<br>(10.2-10.9) | 9.4<br>(9.1-9.6) | 8.1<br>(7.8-8.5) | 7.2<br>(6.9-7.4) |
| **1d** (I) | 10.6<br>(10.3-10.9) | 9.0<br>(8.8-9.3) | 8.0<br>(7.6-8.4) | 6.6<br>(6.4-6.9) |
| **1e** (Me) | 10<br>(9.7-10.3) | 8.3<br>(8.1-8.5) | 8.1<br>(7.7-8.4) | 6.1<br>(5.8-6.4) |
| **1f** (OMe) | 9.8<br>(9.5-10.1) | 7.9<br>(7.7-8.1) | 7.4<br>(7.0-7.7) | 5.6<br>(5.3-5.8) |
| **1g** ($CF_3$) | 9.7<br>(9.4-10) | 8.3<br>(8.1-8.6) | 8.1<br>(7.8-8.4) | 6.6<br>(6.4-6.9) |
| **1h** (CN) | 9.3<br>(9.0-9.5) | 7.4<br>(7.2-7.7) | 7.5<br>(7.2-7.7) | 5.9<br>(5.6-6.1) |
| **1i** ($NO_2$) | 9.0<br>(8.8-9.3) | 7.2<br>(6.9-7.4) | 7.2<br>(6.9-7.5) | 5.3<br>(5.0-5.5) |
| **1j** (2-pyr) | 8.2<br>(8.0-8.5) | 6.6<br>(6.3-6.9) | 7.4<br>(7.1-7.7) | 5.3<br>(5.0-5.6) |
| **1k** (3-pyr) | 7.3<br>(7.0-7.6) | 5.4<br>(5.1-5.8) | 6.6<br>(6.3-6.9) | 3.2<br>(2.7-3.6) |
| **1l** (4-pyr) | 7.5<br>(7.3-7.8) | 6.2<br>(5.9-6.4) | 6.6<br>(6.3-6.8) | 3.9<br>(3.6-4.2) |
| DADLE | 9.0<br>(8.8-9.3) | 8.3<br>(8.0-8.6) | 7.6<br>(7.3-7.9) | 6.4<br>(6.3-6.5) |
| Leu-enkephalin | 8.9<br>(8.7-9.2) | 7.8<br>(7.7-8.0) | 7.0<br>(6.8-7.3) | 5.5<br>(5.2-5.7) |
| DAMGO | 5.9<br>(5.6-6.2) | - | 7.8<br>(7.6-8.0) | 6.9<br>(6.8-7.1) |

**SI-3 Characterization of Peptides**

**$H_2$N-Tyr-Gly-Gly-(*meta*-F)Phe-Leu-OH (1a).** Yield 52.6%, 60.0 mg colorless solid. $^1$H NMR (500 MHz, MeOD-$d_4$) δ 7.29 (q, $J$ = 7.8 Hz, 1H), 7.16 – 7.06 (m, 4H), 6.95 (t, $J$ = 8.6 Hz, 1H), 6.81 (d, $J$ = 8.4 Hz, 2H), 4.74 (dd, $J$ = 9.8, 4.6 Hz, 1H), 4.38 (t, $J$ = 7.3 Hz, 1H), 4.15 – 4.09 (m, 1H), 4.00 – 3.91 (m, 2H), 3.80 – 3.72 (m, 2H), 3.24 – 3.14 (m, 2H), 3.03 – 2.98 (m, 2H), 1.77 – 1.61 (m, 3H), 0.96 (d, $J$ = 6.2 Hz, 3H), 0.92 (d, $J$ = 6.2 Hz, 3H). $^{13}$C NMR (126 MHz, MeOD-$d_4$) δ 173.9, 171.9, 170.1, 169.8, 169.7, 162.8 (d, $J$ = 244.0 Hz), 156.9, 139.8 (d, $J$ = 7.6 Hz), 130.1, 129.6 (d, $J$ = 8.2 Hz), 124.9 (d, $J$ = 2.8 Hz), 124.6, 115.7 (d, $J$ = 21.6 Hz), 115.5, 113.0 (d, $J$ = 21.3 Hz), 54.7, 54.0, 51.4, 42.6, 41.8, 40.3, 37.2 (d, $J$ = 1.8 Hz), 36.3, 24.5, 22.0, 20.5. $^{19}$F NMR (471 MHz, MeOD-$d_4$) δ –115.0. HRMS (ESI$^+$) mass calculated for $[M+H]^+$ ($C_{28}H_{36}FN_5O_7$) m/z 574.2677, found m/z 574.2669; purity ≥95%, rt 2.76 min (protocol A).

**$H_2$N-Tyr-Gly-Gly-(*meta*-Cl)Phe-Leu-OH (1b)**. Yield 50.9%, 60.2 mg colorless solid. $^1$H NMR (500 MHz, MeOD-$d_4$) δ 7.37 (s, 1H), 7.31 – 7.20 (m, 3H), 7.13 (d, $J$ = 8.4 Hz, 2H), 6.81 (d, $J$ = 8.5 Hz, 2H), 4.72 (dd, $J$ = 9.8, 4.6 Hz, 1H), 4.39 (t, $J$ = 7.4 Hz, 1H), 4.18 – 4.06 (m, 1H), 4.00 – 4.10 (m, 2H), 3.80 – 3.73 (m, 2H), 3.78 (d, $J$ = 22.5 Hz, 1H), 3.75 (d, $J$ = 23.1 Hz, 1H), 3.21 – 3.16 (m, 2H), 3.02 – 2.96 (m, 2H), 1.76 – 1.59 (m, 3H), 0.97 (d, $J$ = 6.2 Hz, 3H), 0.92 (d, $J$ = 6.1 Hz, 3H). $^{13}$C NMR (126 MHz, MeOD-$d_4$) δ 174.8, 171.9, 170.0, 169.8, 169.7, 156.9, 139.3, 133.7, 130.1, 129.5, 129.0, 127.5, 126.5, 124.5, 115.5, 54.7, 54.0, 51.3, 42.6, 41.8, 40.3, 37.1, 36.3, 24.5, 22.0, 20.4. HRMS (ESI$^+$) mass calculated for $[M+H]^+$ ($C_{28}H_{36}ClN_5O_7$) m/z 590.2473, found m/z 590.2431; purity ≥95%, rt 1.65 min (protocol B).

**$H_2$N-Tyr-Gly-Gly-(*meta*-Br)Phe-Leu-OH (1c).** Yield 47.6%, 60.5 mg colorless solid. $^1$H NMR (500 MHz, MeOD-$d_4$) δ 7.54 (s, 1H), 7.38 (d, $J$ = 7.9 Hz, 1H), 7.29 (d, $J$ = 7.6 Hz, 1H), 7.21 (t, $J$ = 7.8 Hz, 1H), 7.14 (d, $J$ = 8.2 Hz, 2H), 6.81 (d, $J$ = 8.2 Hz, 2H), 4.71 (dd, $J$ = 10.1, 4.4 Hz, 1H), 4.33 (t, $J$ = 7.3 Hz, 1H), 4.12 (t, $J$ = 7.3 Hz, 1H), 3.99 (d, $J$ = 26.6 Hz, 1H), 3.95 (d, $J$ = 27.1 Hz, 1H), 3.77 (d, $J$ = 24.5 Hz, 1H), 3.73 (d, $J$ = 25.0 Hz, 1H), 3.19 (ddd, $J$ = 15.3, 10.5, 5.4 Hz, 2H), 3.24 – 2.94 (m, 2H), 1.83 – 1.55 (m, 3H), 0.96 (d, $J$ = 6.2 Hz, 3H), 0.92 (d, $J$ = 6.1 Hz, 3H). $^{13}$C NMR (126 MHz, MeOD-$d_4$) δ 171.8, 170.0, 170.0, 169.8, 156.9, 139.7, 132.0, 130.1, 129.7, 129.4, 127.9, 124.6, 121.9, 115.5, 54.8, 54.1, 52.0, 48.1, 47.9, 42.7, 41.8, 40.5, 37.0, 36.4, 24.6,

22.0, 20.5. HRMS ($ESI^+$) mass calculated for $[M+H]^+$ ($C_{28}H_{36}BrN_5O_7$) m/z 656.1696, found m/z 656.1711; purity ≥95%, rt 2.8 min (protocol A).

**$H_2N$-Tyr-Gly-Gly-(*meta*-I)Phe-Leu-OH (1d).** Yield 55.2%, 75.0 mg colorless solid. $^1H$ NMR (500 MHz, MeOD-$d_4$) δ 7.75 (s, 1H), 7.58 (d, *J* = 7.6 Hz, 1H), 7.33 (d, *J* = 7.7 Hz, 1H), 7.14 (d, *J* = 8.1 Hz, 2H), 7.07 (t, *J* = 7.8 Hz, 1H), 6.81 (d, *J* = 8.1 Hz, 2H), 4.68 (dd, *J* = 10.5, 4.1 Hz, 1H), 4.25 (dd, *J* = 9.1, 5.3 Hz, 1H), 4.12 (t, *J* = 7.2 Hz, 1H), 4.03 – 3.94 (m, 2H), 3.77 – 3.67 (m, 2H), 3.74 (d, *J* = 31.4 Hz, 1H), 3.70 (d, *J* = 32.0 Hz, 1H), 3.23 – 3.13 (m, 2H), 2.97 (ddd, *J* = 21.8, 14.0, 9.4 Hz, 2H), 1.76 – 1.54 (m, 3H), 0.96 (d, *J* = 6.1 Hz, 3H), 0.92 (d, *J* = 6.0 Hz, 3H). $^{13}C$ NMR (126 MHz, MeOD-$d_4$) δ 174.3, 171.7, 170.5, 170.1, 169.8, 156.9, 139.9, 138.0, 135.5, 130.1, 129.8, 128.4, 124.7, 115.5, 93.5, 54.8, 54.3, 53.2, 42.9, 41.8, 40.9, 36.9, 36.5, 24.7, 22.2, 20.7. HRMS ($ESI^+$) mass calculated for $[M+H]^+$ ($C_{28}H_{36}IN_5O_7$) m/z 682.1737, found m/z 681.1741; purity ≥95%, rt 2.97 min (protocol A).

**$H_2N$-Tyr-Gly-Gly-(*meta*-Me)Phe-Leu-OH (1e).** Yield 78.9%, 90.2 mg colorless solid. $^1H$ NMR (500 MHz, MeOD-$d_4$) δ 7.20 – 7.11 (m, 4H), 7.08 (d, *J* = 7.7 Hz, 1H), 7.03 (d, *J* = 7.5 Hz, 1H), 6.81 (d, *J* = 8.2 Hz, 2H), 4.69 (dd, *J* = 9.8, 4.6 Hz, 1H), 4.37 (t, *J* = 7.3 Hz, 1H), 4.12 (t, *J* = 7.3 Hz, 1H), 4.02 – 3.88 (m, 2H), 3.77 (d, *J* = 27.8 Hz, 1H), 3.74 (d, *J* = 28.2 Hz, 1H), 3.17 (ddd, *J* = 14.4, 9.3, 5.5 Hz, 2H), 2.97 (ddd, *J* = 23.4, 14.0, 9.0 Hz, 2H), 1.73 – 1.63 (m, 3H), 0.96 (d, *J* = 6.3 Hz, 3H), 0.92 (d, *J* = 6.2 Hz, 3H). $^{13}C$ NMR (126 MHz, MeOD-$d_4$) δ 175.2, 172.2, 170.0, 169.8, 169.7, 156.9, 137.7, 136.8, 130.1, 129.6, 127.9, 127.0, 126.0, 124.6, 115.4, 54.8, 54.5, 51.6, 42.6, 41.8, 40.4, 37.4, 36.3, 24.5, 22.0, 20.5, 20.0. HRMS ($ESI^+$) mass calculated for $[M+H]^+$ ($C_{29}H_{39}N_5O_7$) m/z 570.2927, found m/z 570.2923; purity ≥95%, rt 2.88 min (protocol A).

**$H_2N$-Tyr-Gly-Gly-(*meta*-OMe)Phe-Leu-OH (1f).** Yield 22.3%, 26.0 mg colorless solid. $^1H$ NMR (500 MHz, MeOD-$d_4$) δ 7.19 (t, *J* = 7.9 Hz, 1H), 7.13 (d, *J* = 8.3 Hz, 2H), 6.91 – 6.85 (m, 2H), 6.84 – 6.74 (m, 3H), 4.73 (dd, *J* = 9.6, 4.7 Hz, 1H), 4.43 (t, *J* = 7.4 Hz, 1H), 4.13 – 4.08 (m, 1H), 4.00 – 3.92 (m, 2H), 3.83 – 3.71 (m, 5H), 3.20-3.15 (m, 2H), 3.06 – 2.91 (m, 2H), 1.75 – 1.62 (m, 3H), 0.97 (d, *J* = 6.3 Hz, 3H), 0.92 (d, *J* = 6.4 Hz, 3H). $^{13}C$ NMR (126 MHz, MeOD-$d_4$) δ 174.2, 172.3, 170.0, 169.7, 169.6, 159.8, 156.9, 138.4, 130.1, 128.9, 124.5, 121.3, 115.4, 114.3, 112.1, 54.7, 54.2, 54.2, 50.7, 42.5, 41.8, 40.1, 37.5, 36.2, 24.5, 21.9, 20.4. HRMS ($ESI^+$) mass calculated for $[M+H]^+$ ($C_{29}H_{39}N_5O_8$) m/z 586.2877, found m/z 585.2932; purity ≥95%, rt 1.57 min (protocol B).

**$H_2N$-Tyr-Gly-Gly-(*meta*-$CF_3$)Phe-Leu-OH (1g).** Yield 32.1%, 40.1 mg colorless solid. $^1H$ NMR (500 MHz, MeOD-$d_4$) δ 7.67 (s, 1H), 7.60 – 7.46 (m, 3H), 7.13 (d, *J* = 8.2 Hz, 2H), 6.80 (d, *J* = 8.2 Hz, 2H), 4.76 (dd, *J* = 9.8, 4.7 Hz, 1H), 4.41 (t, *J* = 7.4 Hz, 1H), 4.14 – 4.11 (m, 1H), 3.95 (t, *J* = 16.2 Hz, 1H), 3.76 (dd, *J* = 32.2, 16.8 Hz, 1H), 3.29 (dd, *J* = 14.0, 4.7 Hz, 1H), 3.18 (dd, *J* = 14.2, 6.5 Hz, 1H), 3.08 (dd, *J* = 14.0, 9.8 Hz, 1H), 3.00 (dd, *J* = 14.2, 8.1 Hz, 1H), 1.77 – 1.62 (m, 3H), 0.97 (d, *J* = 6.5 Hz, 1H), 0.92 (d, *J* = 6.2 Hz, 1H). $^{13}C$ NMR (126 MHz, MeOD-$d_4$) δ 174.5, 171.9, 170.0, 169.8, 169.7, 156.9, 138.4, 132.9, 130.2 (d, *J* = 31.8 Hz), 130.1, 128.7, 125.7 (q, *J* = 3.7 Hz), 124.5, 124.3 (q, *J* = 272.2 Hz), 123.1 (q, *J* = 4.0 Hz), 115.4, 54.7, 54.0, 51.0, 42.6, 41.7, 40.2, 37.3, 36.3, 24.5, 21.9, 20.4. $^{19}F$ NMR (471 MHz, MeOD-$d_4$) δ –63.3. HRMS ($ESI^+$) mass calculated for $[M+H]^+$ ($C_{29}H_{36}F_3N_5O_7$) m/z 624.2645, found m/z 624.2618; purity ≥95%, rt 2.89 min (protocol A).

**$H_2N$-Tyr-Gly-Gly-(*meta*-CN)Phe-Leu-OH (1h).** Yield 43.1%, 50.0 mg colorless solid. $^1H$ NMR (500 MHz, MeOD-$d_4$) δ 7.70 (s, 1H), 7.63 – 7.59 (m, 2H), 7.48 (t, *J* = 7.7 Hz, 1H), 7.14 (d, *J* = 8.0 Hz, 2H), 6.81 (d, *J* = 8.0 Hz, 2H), 4.75 (dd, *J* = 9.6, 4.8 Hz, 1H), 4.38 (dd, *J* = 9.0, 5.4 Hz, 1H), 4.13 (t, *J* = 7.4 Hz, 1H), 4.00-3.91 (m, 2H), 3.81-3.71 (m, 2H), 3.26 (dd, *J* = 14.0, 4.8 Hz, 1H), 3.19 (dd, *J* = 14.2, 6.4 Hz, 1H), 3.08-2.99 (m, 2H), 1.78 – 1.58 (m, 3H), 0.97 (d, *J* = 6.1 Hz, 3H), 0.92 (d, *J* = 6.0 Hz, 3H). $^{13}C$ NMR (126 MHz, MeOD-$d_4$) δ 173.9, 171.6, 170.0, 169.9, 169.7, 156.9, 138.8, 134.0, 132.8, 130.2, 130.1, 129.1, 124.6, 118.5, 115.5, 111.9, 54.8, 53.8, 51.5, 42.7, 41.8, 40.3, 37.0, 36.3, 24.6, 22.0, 20.4. HRMS ($ESI^+$) mass calculated for $[M+H]^+$ ($C_{29}H_{36}N_6O_7$) m/z 581.2723, found m/z 581.2755; purity ≥95%, rt 1.56 min (protocol B).

**$H_2N$-Tyr-Gly-Gly-(*meta*-$NO_2$)Phe-Leu-OH (1i).** Yield 66.7%, 80.3 mg colorless solid. $^1H$ NMR (400 MHz, MeOD-$d_4$) δ 8.30 (s, 1H), 8.11 (d, *J* = 8.2 Hz, 1H), 7.73 (d, *J* = 7.7 Hz, 1H), 7.54 (t, *J* = 7.9 Hz, 1H), 7.15 (d, *J* = 8.5 Hz, 2H), 6.81 (d, *J* = 8.5 Hz, 2H), 4.77 (dd, *J* = 10.6, 4.2 Hz, 1H), 4.25 (dd, *J* = 8.9, 5.5 Hz, 1H), 4.15 (dd, *J* = 8.3, 6.2 Hz, 1H), 3.96 (dd, *J* = 29.6, 16.7 Hz, 2H), 3.69 (dd, *J* = 45.1, 16.8 Hz, 2H), 3.37 (dd, *J* = 13.9, 4.3 Hz, 1H), 3.24 – 3.10 (m, 2H), 3.01 (dd, *J* = 14.2, 8.3 Hz, 1H), 1.73 – 160 (m, 3H), 0.96 (d, *J* = 6.1 Hz, 3H), 0.92 (d, *J* = 6.0 Hz, 3H). $^{13}C$ NMR (126 MHz, MeOD-$d_4$) δ 177.2, 171.4, 170.6, 170.0, 169.8, 156.9, 148.2, 139.5, 135.5, 130.1, 129.1, 124.7, 124.1, 121.3, 115.5, 54.9, 54.0, 53.2, 43.0, 41.8, 40.9, 37.0, 36.5,

24.7, 22.1, 20.6. HRMS (ESI$^+$) mass calculated for $[M+H]^+$ ($C_{29}H_{36}N_6O_9$) m/z 601.2606, found m/z 601.2622; purity ≥95%, rt 1.59 min (protocol B).

**$H_2$N-Tyr-Gly-Gly-(2-pyridyl)Ala-Leu-OH (1j).** Yield 10.8%, 12.1 mg colorless solid. $^1$H NMR (500 MHz, MeOD-$d_4$) δ 8.63 (d, *J* = 5.4 Hz, 1H), 8.18 (t, *J* = 7.8 Hz, 1H), 7.73 (d, *J* = 7.9 Hz, 1H), 7.68 – 7.61 (m, 1H), 7.13 (d, *J* = 8.3 Hz, 2H), 6.80 (d, *J* = 8.2 Hz, 2H), 4.93 (dd, *J* = 8.5, 5.5 Hz, 1H), 4.44 – 4.38 (m, 1H), 4.11 (t, *J* = 7.2 Hz, 1H), 4.00 – 3.76 (m, 4H), 3.50 (dd, *J* = 14.3, 5.4 Hz, 1H), 3.38 – 3.27 (m, 1H), 3.17 (dd, *J* = 14.2, 6.5 Hz, 1H), 3.01 (dd, *J* = 14.2, 8.0 Hz, 1H), 1.78 – 1.58 (m, 3H), 0.97 (d, *J* = 6.0 Hz, 3H), 0.92 (d, *J* = 6.0 Hz, 3H). $^{13}$C NMR (126 MHz, MeOD-$d_4$) δ 174.1, 170.8, 170.2, 169.8, 169.6, 156.9, 154.5, 144.7, 142.1, 130.1, 126.5, 124.5, 123.8, 115.4, 54.7, 52.4, 50.9, 42.4, 41.9, 39.9, 37.4, 36.2, 24.5, 21.9, 20.3. HRMS (ESI$^+$) mass calculated for $[M+H]^+$ ($C_{27}H_{36}N_6O_7$) m/z 557.2723, found m/z 557.2748; purity ≥95%, rt 2.46 min (protocol A).

**$H_2$N-Tyr-Gly-Gly-(3-pyridyl)Ala-Leu-OH (1k).** Yield 62.9%, 70.0 mg colorless solid. $^1$H NMR (400 MHz, MeOD-$d_4$) δ 8.49 (s, 1H), 8.42 (d, *J* = 3.8 Hz, 1H), 7.83 (d, *J* = 7.9 Hz, 1H), 7.40 (dd, *J* = 7.8, 5.0 Hz, 1H), 7.14 (d, *J* = 8.5 Hz, 2H), 6.81 (d, *J* = 8.5 Hz, 2H), 4.77 (dd, *J* = 9.3, 4.9 Hz, 1H), 4.45 – 4.33 (m, 1H), 4.12 (dd, *J* = 8.2, 6.4 Hz, 1H), 3.95 (dd, *J* = 27.3, 16.7 Hz, 2H), 3.78 (dd, *J* = 19.3, 16.7 Hz, 2H), 3.28 – 3.16 (m, 2H), 3.09 – 2.97 (m, 2H), 1.77 – 1.57 (m, 3H), 0.97 (d, *J* = 6.2 Hz, 3H), 0.92 (d, *J* = 6.1 Hz, 3H). $^{13}$C NMR (126 MHz, MeOD-$d_4$) δ 174.6, 173.8, 171.5, 170.1, 169.7, 156.9, 149.2, 146.7, 138.3, 133.6, 130.1, 124.5, 123.8, 115.5, 54.7, 53.5, 51.1, 42.5, 41.8, 40.2, 36.3, 34.7, 24.6, 21.9, 20.4. HRMS (ESI$^+$) mass calculated for $[M+H]^+$ ($C_{27}H_{36}N_6O_7$) m/z 557.2723, found m/z 557.2659; purity ≥95%, rt 2.46 min (protocol A).

**$H_2$N-Tyr-Gly-Gly-(4-pyridyl)Ala-Leu-OH (1l).** Yield 32.4%, 36.1 mg colorless solid. $^1$H NMR (500 MHz, MeOD-$d_4$) δ 8.51 (d, *J* = 5.3 Hz, 2H), 7.53 (d, *J* = 5.3 Hz, 2H), 7.13 (d, *J* = 8.2 Hz, 2H), 6.80 (d, *J* = 8.3 Hz, 2H), 4.84 (dd, *J* = 9.4, 4.9 Hz, 1H), 4.45 – 4.36 (m, 1H), 4.13 (dd, *J* = 8.1, 6.4 Hz, 1H), 3.94 (dd, *J* = 20.4, 16.7 Hz, 2H), 3.78 (dd, *J* = 30.3, 16.7 Hz, 2H), 3.33 – 3.27 (m, 1H), 3.21 – 3.08 (m, 2H), 3.01 (dd, *J* = 14.2, 8.1 Hz, 1H), 1.75 – 1.63 (m, 3H), 0.97 (d, *J* = 6.1 Hz, 3H), 0.92 (d, *J* = 6.1 Hz, 3H). $^{13}$C NMR (126 MHz, MeOD-$d_4$) δ 174.3, 173.8, 171.2, 170.1, 169.7, 156.9, 150.2, 146.7, 130.1, 125.8, 124.5, 115.4, 54.7, 52.9, 50.9, 42.6, 41.8, 40.0,

37.1, 36.2, 24.5, 21.9, 20.3. HRMS ($ESI^+$) mass calculated for $[M+H]^+$ ($C_{27}H_{36}N_6O_7$) m/z 557.2723, found m/z 557.2729; purity ≥95%, rt 1.20 min (protocol B).

### NMR Spectra of Peptides

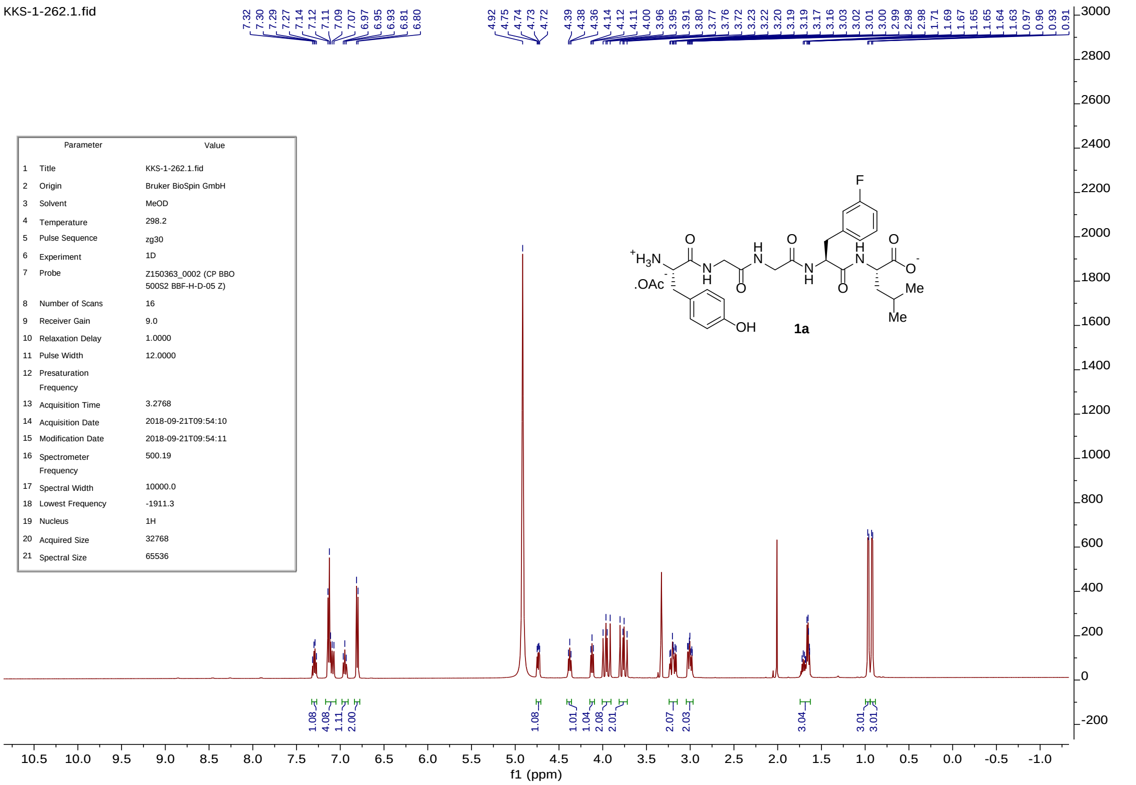


| | Parameter | Value |
|---|---|---|
| 1 | Title | KKS-1-262.1.fid |
| 2 | Origin | Bruker BioSpin GmbH |
| 3 | Solvent | MeOD |
| 4 | Temperature | 298.2 |
| 5 | Pulse Sequence | zg30 |
| 6 | Experiment | 1D |
| 7 | Probe | Z150363_0002 (CP BBO 500S2 BBF-H-D-05 Z) |
| 8 | Number of Scans | 16 |
| 9 | Receiver Gain | 9.0 |
| 10 | Relaxation Delay | 1.0000 |
| 11 | Pulse Width | 12.0000 |
| 12 | Presaturation Frequency | |
| 13 | Acquisition Time | 3.2768 |
| 14 | Acquisition Date | 2018-09-21T09:54:10 |
| 15 | Modification Date | 2018-09-21T09:54:11 |
| 16 | Spectrometer Frequency | 500.19 |
| 17 | Spectral Width | 10000.0 |
| 18 | Lowest Frequency | -1911.3 |
| 19 | Nucleus | 1H |
| 20 | Acquired Size | 32768 |
| 21 | Spectral Size | 65536 |

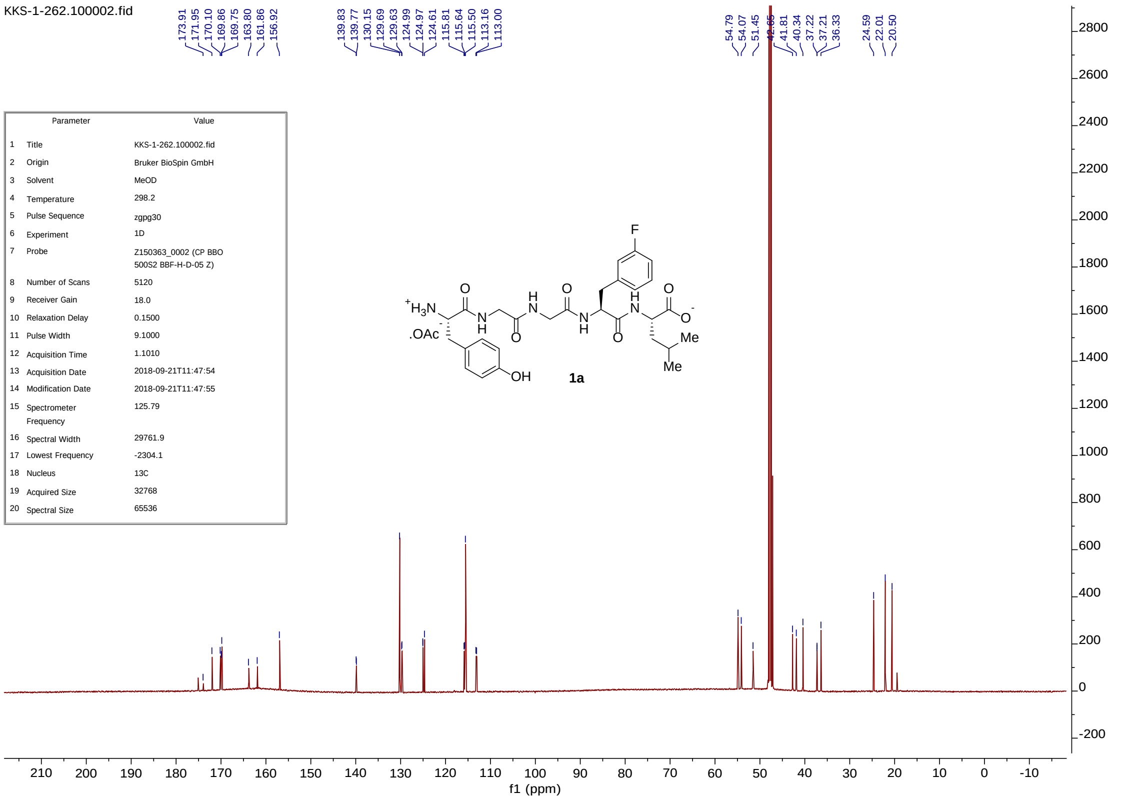


| | Parameter | Value |
|---|---|---|
| 1 | Title | KKS-1-262.100002.fid |
| 2 | Origin | Bruker BioSpin GmbH |
| 3 | Solvent | MeOD |
| 4 | Temperature | 298.2 |
| 5 | Pulse Sequence | zgpg30 |
| 6 | Experiment | 1D |
| 7 | Probe | Z150363_0002 (CP BBO 500S2 BBF-H-D-05 Z) |
| 8 | Number of Scans | 5120 |
| 9 | Receiver Gain | 18.0 |
| 10 | Relaxation Delay | 0.1500 |
| 11 | Pulse Width | 9.1000 |
| 12 | Acquisition Time | 1.1010 |
| 13 | Acquisition Date | 2018-09-21T11:47:54 |
| 14 | Modification Date | 2018-09-21T11:47:55 |
| 15 | Spectrometer Frequency | 125.79 |
| 16 | Spectral Width | 29761.9 |
| 17 | Lowest Frequency | -2304.1 |
| 18 | Nucleus | 13C |
| 19 | Acquired Size | 32768 |
| 20 | Spectral Size | 65536 |

kks-1-262_fr4.1.fid

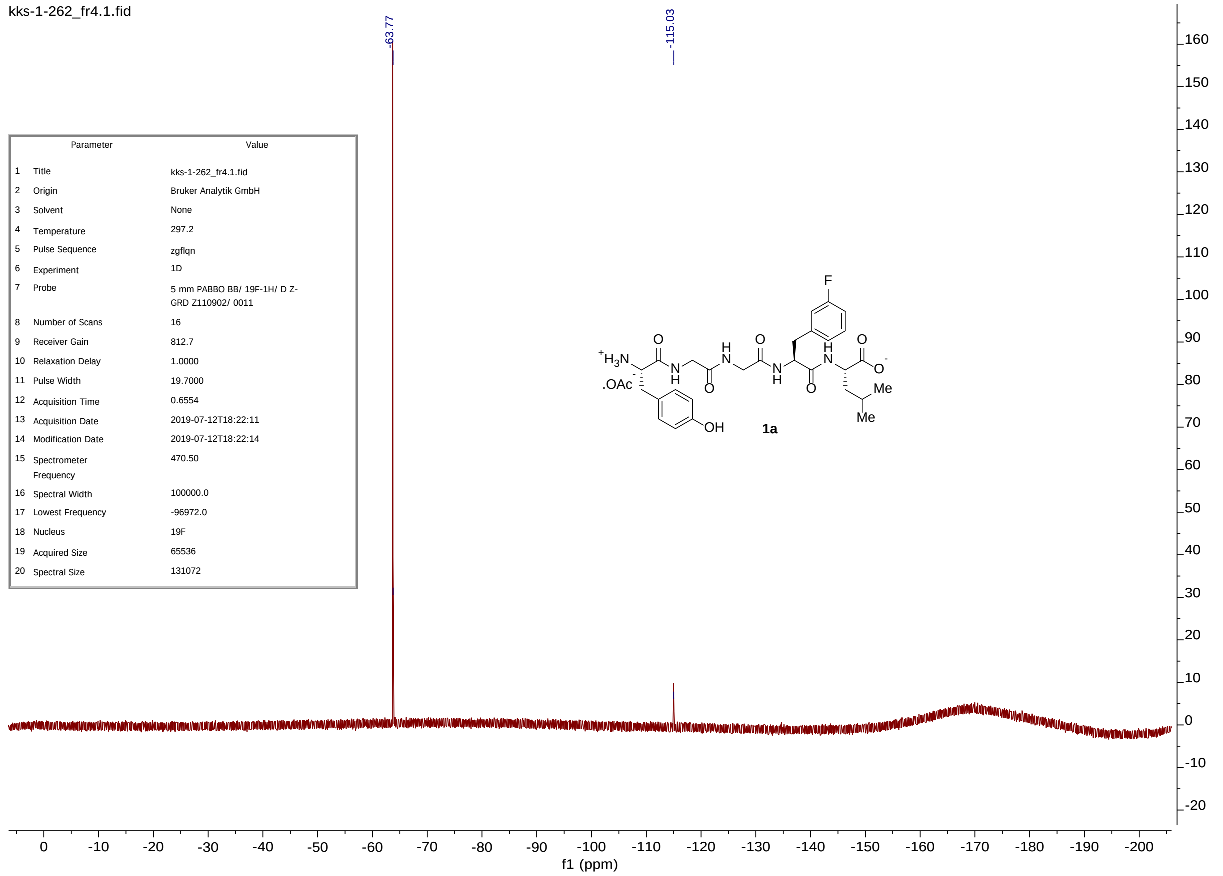


| | Parameter | Value |
|---|---|---|
| 1 | Title | kks-1-262_fr4.1.fid |
| 2 | Origin | Bruker Analytik GmbH |
| 3 | Solvent | None |
| 4 | Temperature | 297.2 |
| 5 | Pulse Sequence | zgflqn |
| 6 | Experiment | 1D |
| 7 | Probe | 5 mm PABBO BB/ 19F-1H/ D Z-GRD Z110902/ 0011 |
| 8 | Number of Scans | 16 |
| 9 | Receiver Gain | 812.7 |
| 10 | Relaxation Delay | 1.0000 |
| 11 | Pulse Width | 19.7000 |
| 12 | Acquisition Time | 0.6554 |
| 13 | Acquisition Date | 2019-07-12T18:22:11 |
| 14 | Modification Date | 2019-07-12T18:22:14 |
| 15 | Spectrometer Frequency | 470.50 |
| 16 | Spectral Width | 100000.0 |
| 17 | Lowest Frequency | -96972.0 |
| 18 | Nucleus | 19F |
| 19 | Acquired Size | 65536 |
| 20 | Spectral Size | 131072 |

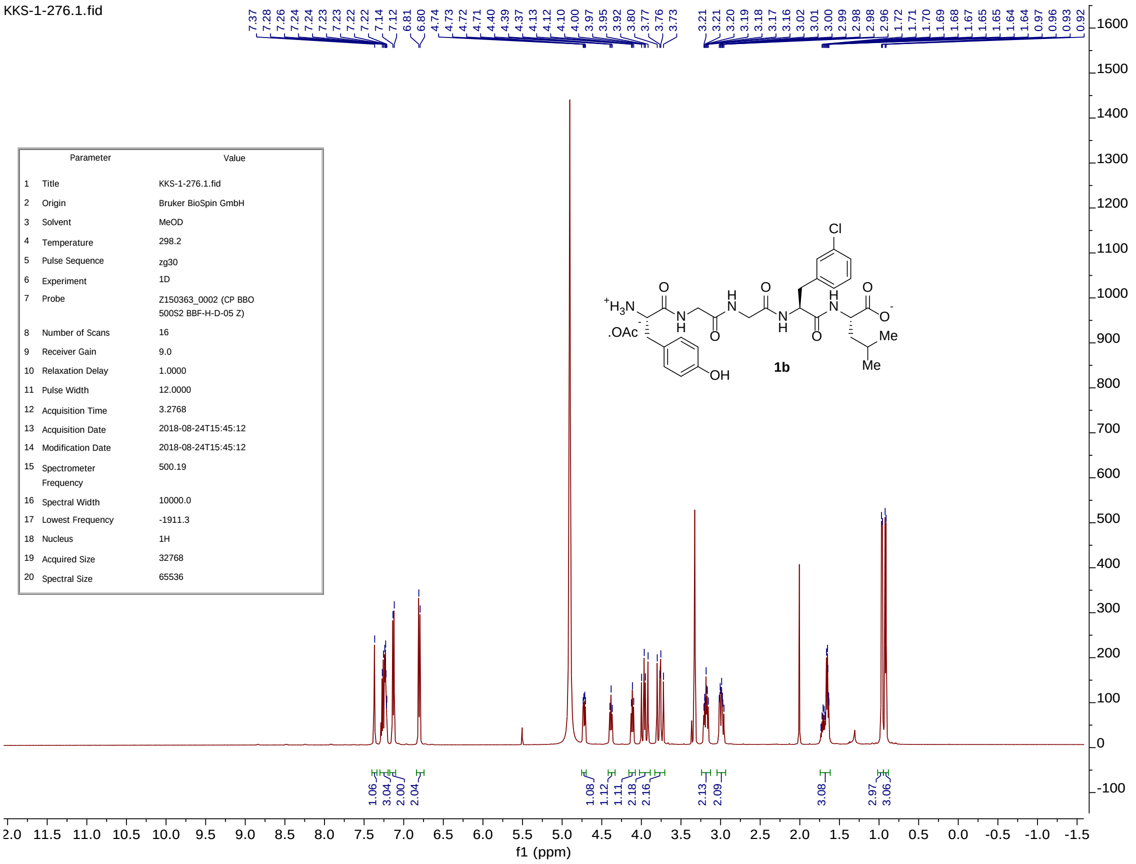

KKS-1-276.1.fid
7.37 7.28 7.26 7.24 7.24 7.23 7.23 7.22 7.22 7.14 7.12 6.81 6.80 4.74 4.73 4.72 4.71 4.40 4.39 4.37 4.13 4.12 4.10 4.00 3.97 3.95 3.92 3.80 3.77 3.76 3.73 3.21 3.21 3.20 3.19 3.18 3.17 3.16 3.02 3.01 3.00 2.99 2.98 2.98 2.96 1.72 1.71 1.70 1.69 1.68 1.67 1.65 1.65 1.64 0.97 0.96 0.93 0.92
Parameter Value
1 Title KKS-1-276.1.fid
2 Origin Bruker BioSpin GmbH
3 Solvent MeOD
4 Temperature 298.2
5 Pulse Sequence zg30
6 Experiment 1D
7 Probe Z150363_0002 (CP BBO 500S2 BBF-H-D-05 Z)
8 Number of Scans 16
9 Receiver Gain 9.0
10 Relaxation Delay 1.0000
11 Pulse Width 12.0000
12 Acquisition Time 3.2768
13 Acquisition Date 2018-08-24T15:45:12
14 Modification Date 2018-08-24T15:45:12
15 Spectrometer Frequency 500.19
16 Spectral Width 10000.0
17 Lowest Frequency -1911.3
18 Nucleus 1H
19 Acquired Size 32768
20 Spectral Size 65536
1b
1.06 3.04 2.00 2.04 1.08 1.12 1.11 2.18 2.16 2.13 2.09 3.08 2.97 3.06
f1 (ppm)

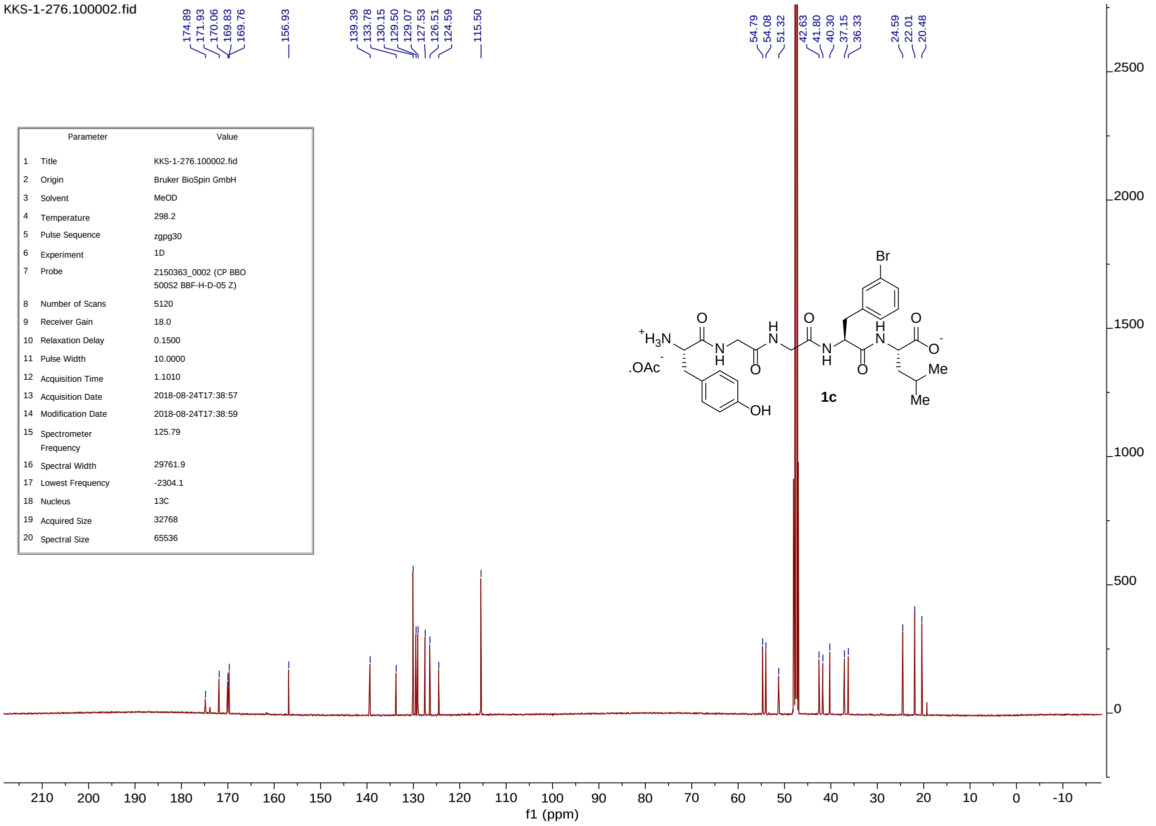
KKS-1-276.100002.fid
174.89
171.93
170.06
169.83
169.76
156.93
139.39
133.78
130.15
129.50
129.07
127.53
126.51
124.59
115.50
54.79
54.08
51.32
42.63
41.80
40.30
37.15
36.33
24.59
22.01
20.48
Parameter Value
1 Title KKS-1-276.100002.fid
2 Origin Bruker BioSpin GmbH
3 Solvent MeOD
4 Temperature 298.2
5 Pulse Sequence zgpg30
6 Experiment 1D
7 Probe Z150363_0002 (CP BBO 500S2 BBF-H-D-05 Z)
8 Number of Scans 5120
9 Receiver Gain 18.0
10 Relaxation Delay 0.1500
11 Pulse Width 10.0000
12 Acquisition Time 1.1010
13 Acquisition Date 2018-08-24T17:38:57
14 Modification Date 2018-08-24T17:38:59
15 Spectrometer Frequency 125.79
16 Spectral Width 29761.9
17 Lowest Frequency -2304.1
18 Nucleus 13C
19 Acquired Size 32768
20 Spectral Size 65536
Br
1c
f1 (ppm)

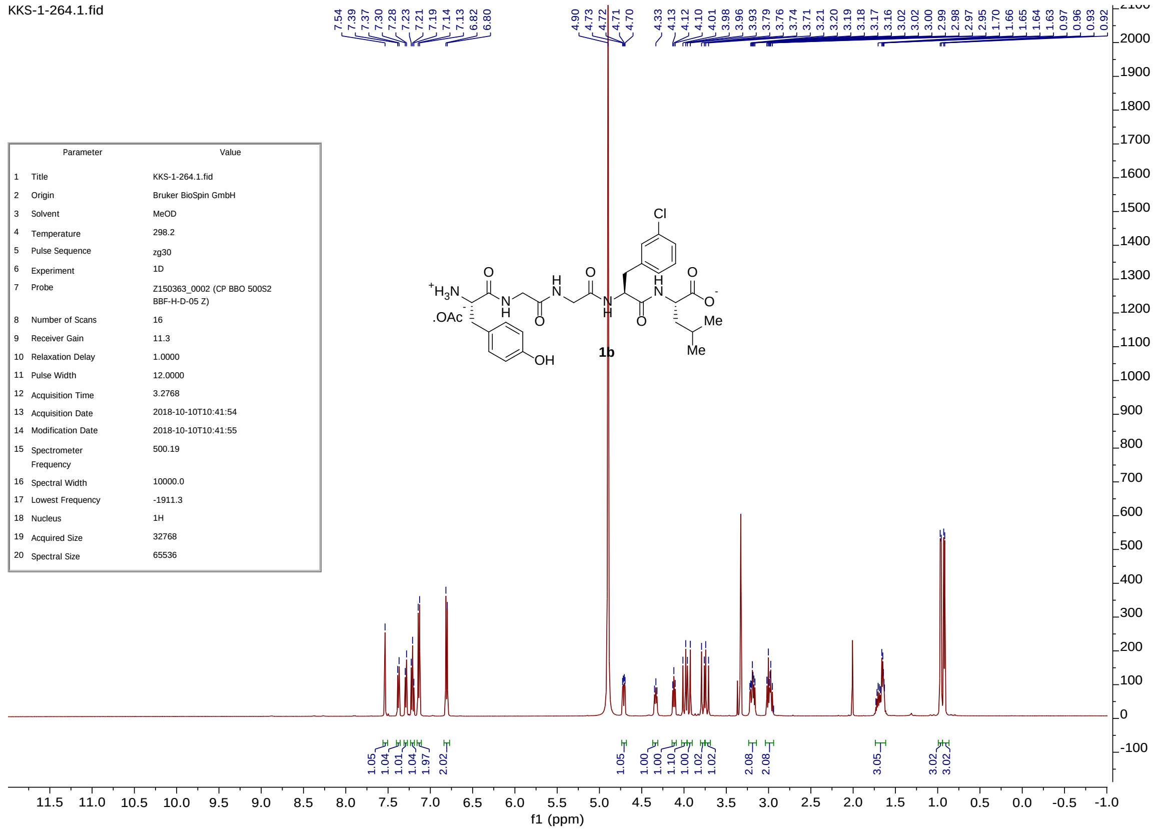
KKS-1-264.1.fid
7.54
7.39
7.37
7.30
7.28
7.23
7.21
7.19
7.14
7.13
6.82
6.80
4.90
4.73
4.72
4.71
4.70
4.33
4.13
4.12
4.10
4.01
3.98
3.96
3.93
3.79
3.76
3.74
3.71
3.21
3.20
3.19
3.18
3.17
3.16
3.02
3.00
2.99
2.97
2.95
1.70
1.66
1.65
1.64
1.63
0.97
0.96
0.93
0.92
Parameter Value
1 Title KKS-1-264.1.fid
2 Origin Bruker BioSpin GmbH
3 Solvent MeOD
4 Temperature 298.2
5 Pulse Sequence zg30
6 Experiment 1D
7 Probe Z150363_0002 (CP BBO 500S2 BBF-H-D-05 Z)
8 Number of Scans 16
9 Receiver Gain 11.3
10 Relaxation Delay 1.0000
11 Pulse Width 12.0000
12 Acquisition Time 3.2768
13 Acquisition Date 2018-10-10T10:41:54
14 Modification Date 2018-10-10T10:41:55
15 Spectrometer Frequency 500.19
16 Spectral Width 10000.0
17 Lowest Frequency -1911.3
18 Nucleus 1H
19 Acquired Size 32768
20 Spectral Size 65536
Cl
1b
f1 (ppm)

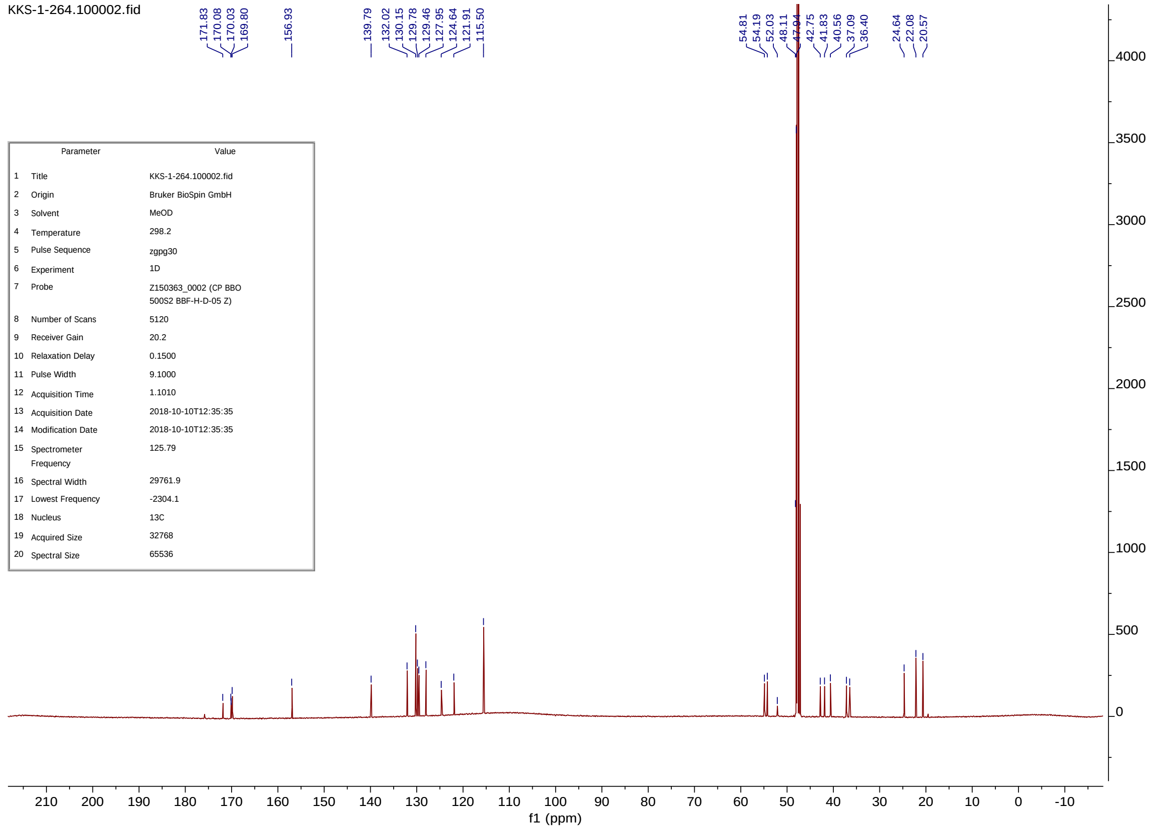


| | Parameter | Value |
|---|---|---|
| 1 | Title | KKS-1-264.100002.fid |
| 2 | Origin | Bruker BioSpin GmbH |
| 3 | Solvent | MeOD |
| 4 | Temperature | 298.2 |
| 5 | Pulse Sequence | zgpg30 |
| 6 | Experiment | 1D |
| 7 | Probe | Z150363_0002 (CP BBO 500S2 BBF-H-D-05 Z) |
| 8 | Number of Scans | 5120 |
| 9 | Receiver Gain | 20.2 |
| 10 | Relaxation Delay | 0.1500 |
| 11 | Pulse Width | 9.1000 |
| 12 | Acquisition Time | 1.1010 |
| 13 | Acquisition Date | 2018-10-10T12:35:35 |
| 14 | Modification Date | 2018-10-10T12:35:35 |
| 15 | Spectrometer Frequency | 125.79 |
| 16 | Spectral Width | 29761.9 |
| 17 | Lowest Frequency | -2304.1 |
| 18 | Nucleus | 13C |
| 19 | Acquired Size | 32768 |
| 20 | Spectral Size | 65536 |

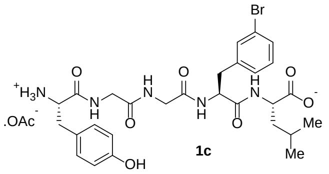


**1c**

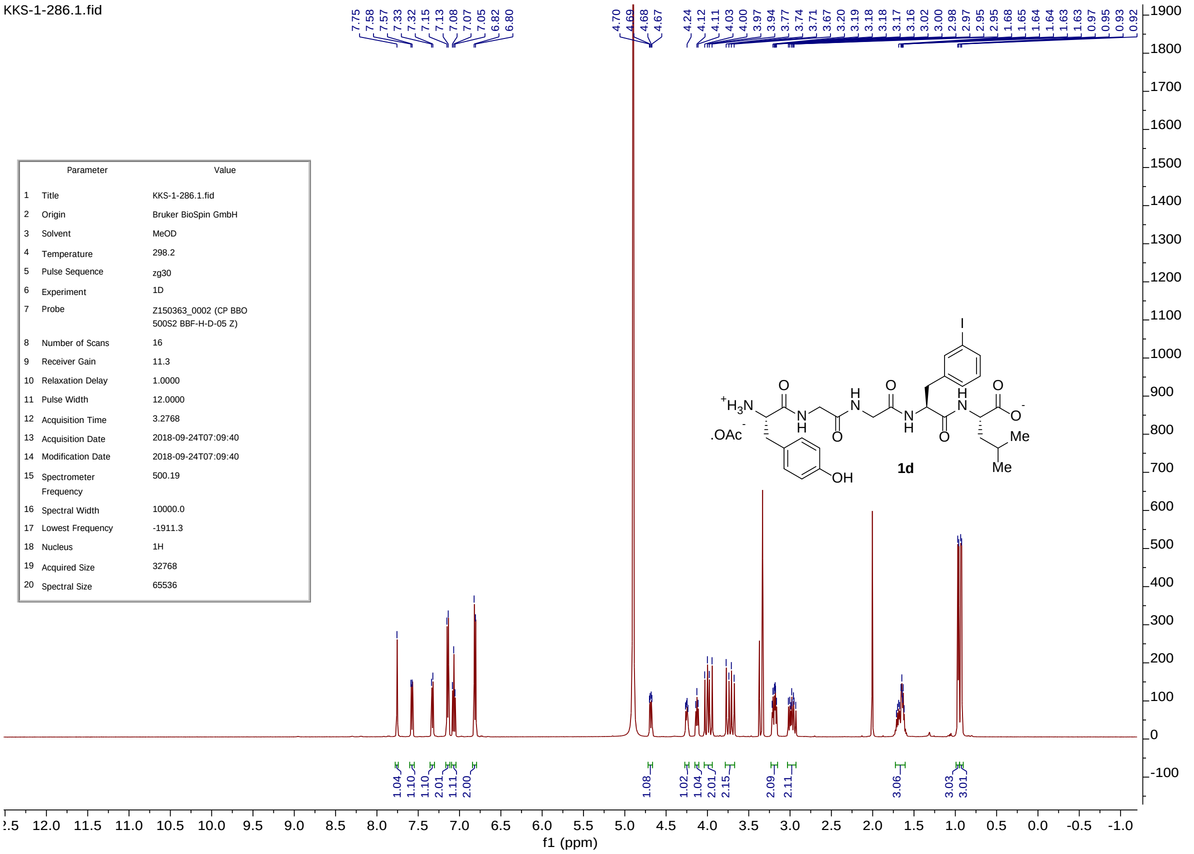


| | Parameter | Value |
|---|---|---|
| 1 | Title | KKS-1-286.1.fid |
| 2 | Origin | Bruker BioSpin GmbH |
| 3 | Solvent | MeOD |
| 4 | Temperature | 298.2 |
| 5 | Pulse Sequence | zg30 |
| 6 | Experiment | 1D |
| 7 | Probe | Z150363_0002 (CP BBO 500S2 BBF-H-D-05 Z) |
| 8 | Number of Scans | 16 |
| 9 | Receiver Gain | 11.3 |
| 10 | Relaxation Delay | 1.0000 |
| 11 | Pulse Width | 12.0000 |
| 12 | Acquisition Time | 3.2768 |
| 13 | Acquisition Date | 2018-09-24T07:09:40 |
| 14 | Modification Date | 2018-09-24T07:09:40 |
| 15 | Spectrometer Frequency | 500.19 |
| 16 | Spectral Width | 10000.0 |
| 17 | Lowest Frequency | -1911.3 |
| 18 | Nucleus | 1H |
| 19 | Acquired Size | 32768 |
| 20 | Spectral Size | 65536 |

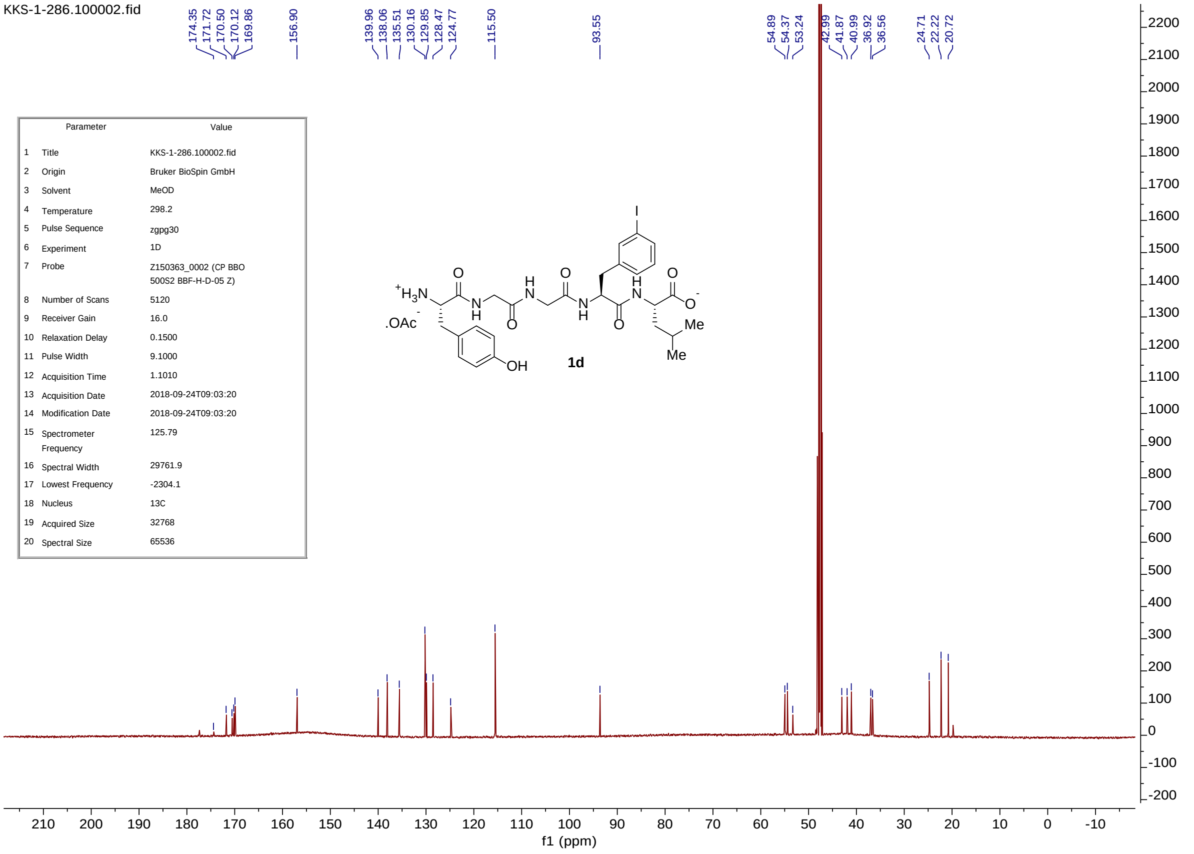


| | Parameter | Value |
|---|---|---|
| 1 | Title | KKS-1-286.100002.fid |
| 2 | Origin | Bruker BioSpin GmbH |
| 3 | Solvent | MeOD |
| 4 | Temperature | 298.2 |
| 5 | Pulse Sequence | zgpg30 |
| 6 | Experiment | 1D |
| 7 | Probe | Z150363_0002 (CP BBO 500S2 BBF-H-D-05 Z) |
| 8 | Number of Scans | 5120 |
| 9 | Receiver Gain | 16.0 |
| 10 | Relaxation Delay | 0.1500 |
| 11 | Pulse Width | 9.1000 |
| 12 | Acquisition Time | 1.1010 |
| 13 | Acquisition Date | 2018-09-24T09:03:20 |
| 14 | Modification Date | 2018-09-24T09:03:20 |
| 15 | Spectrometer Frequency | 125.79 |
| 16 | Spectral Width | 29761.9 |
| 17 | Lowest Frequency | -2304.1 |
| 18 | Nucleus | 13C |
| 19 | Acquired Size | 32768 |
| 20 | Spectral Size | 65536 |

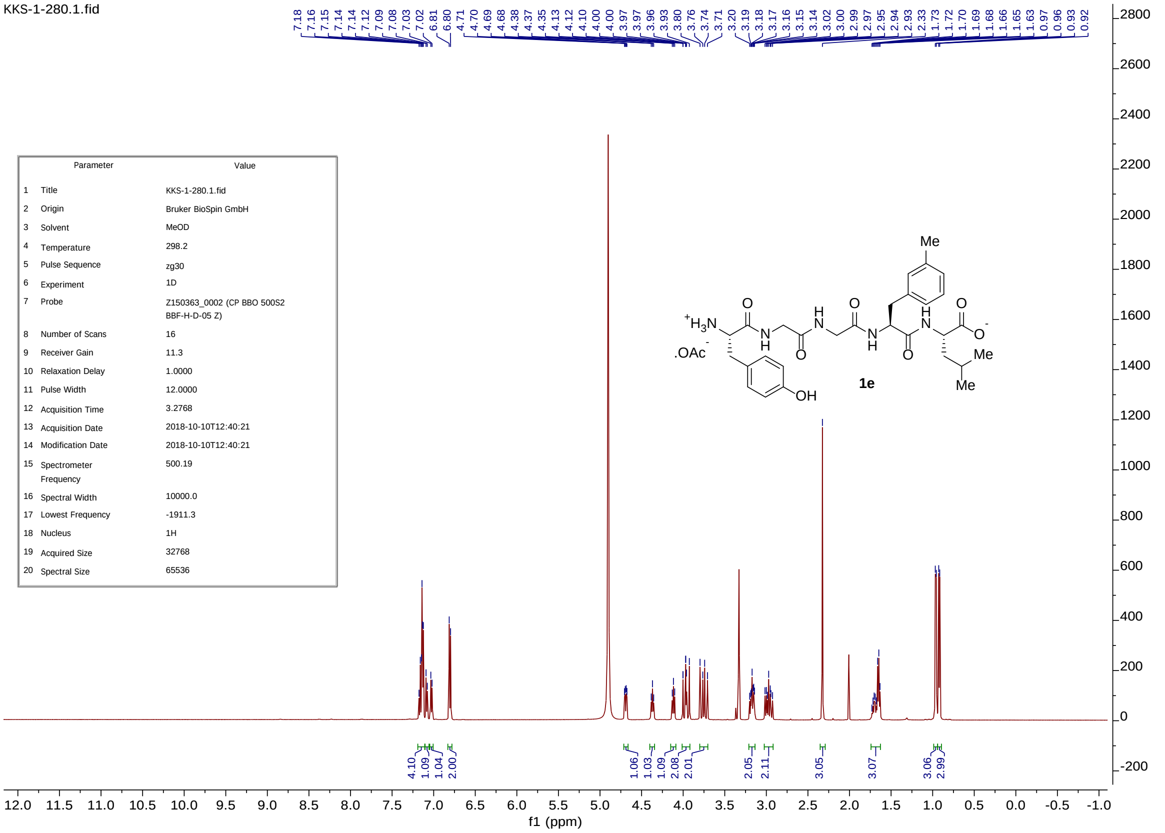
KKS-1-280.1.fid
7.18 7.16 7.15 7.14 7.14 7.12 7.09 7.08 7.03 7.02 6.81 6.80 4.71 4.70 4.69 4.68 4.38 4.37 4.35 4.13 4.12 4.10 4.00 4.00 3.97 3.97 3.96 3.93 3.80 3.76 3.74 3.71 3.20 3.19 3.18 3.17 3.16 3.15 3.14 3.02 3.00 2.99 2.97 2.95 2.94 2.93 2.33 1.73 1.72 1.70 1.69 1.68 1.66 1.65 1.63 0.97 0.96 0.93 0.92
Parameter Value
1 Title KKS-1-280.1.fid
2 Origin Bruker BioSpin GmbH
3 Solvent MeOD
4 Temperature 298.2
5 Pulse Sequence zg30
6 Experiment 1D
7 Probe Z150363_0002 (CP BBO 500S2 BBF-H-D-05 Z)
8 Number of Scans 16
9 Receiver Gain 11.3
10 Relaxation Delay 1.0000
11 Pulse Width 12.0000
12 Acquisition Time 3.2768
13 Acquisition Date 2018-10-10T12:40:21
14 Modification Date 2018-10-10T12:40:21
15 Spectrometer Frequency 500.19
16 Spectral Width 10000.0
17 Lowest Frequency -1911.3
18 Nucleus 1H
19 Acquired Size 32768
20 Spectral Size 65536
Me
$^{+}H_3N$
.OAc$^{-}$
OH
O
N
H
O
O
$O^{-}$
Me
Me
1e
4.10 1.09 1.04 2.00 1.06 1.03 1.09 2.08 2.01 2.05 2.11 3.05 3.07 3.06 2.99
f1 (ppm)

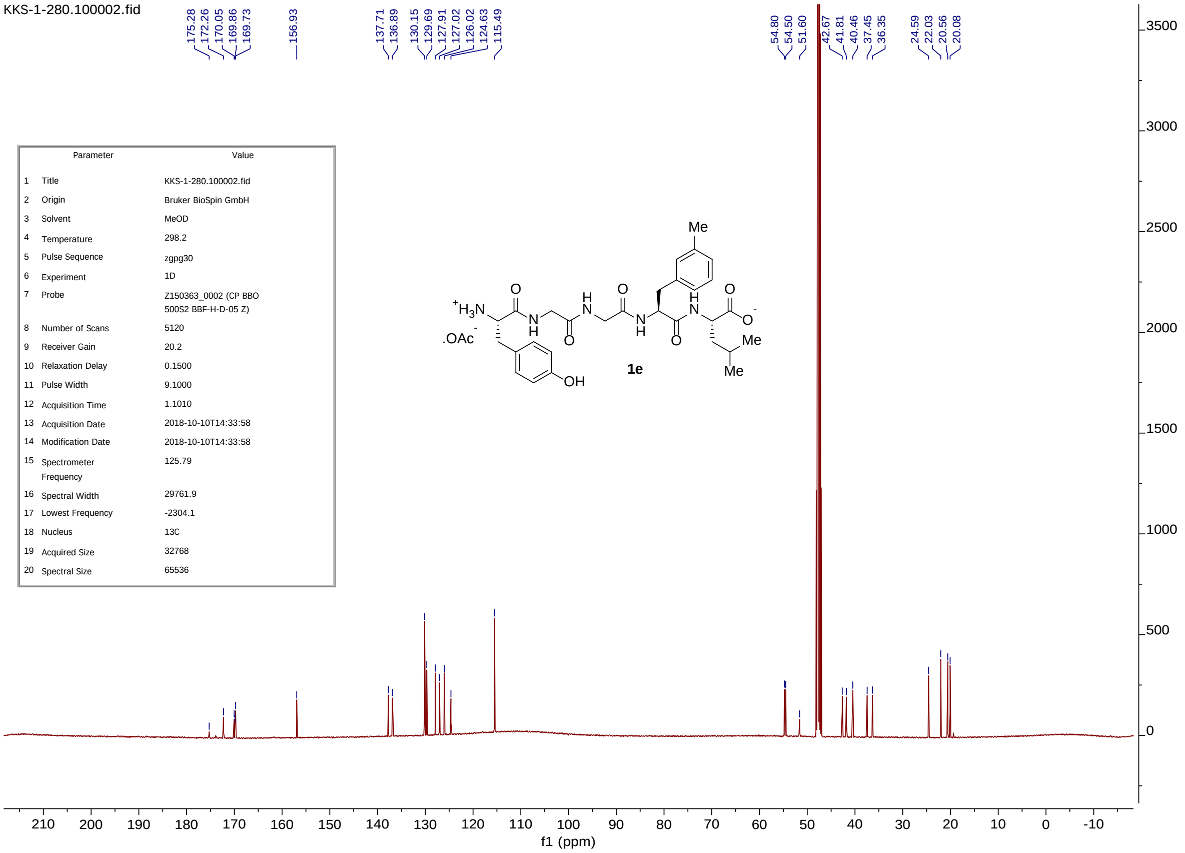
KKS-1-280.100002.fid
175.28 172.26 170.05 169.86 169.73 156.93 137.71 136.89 130.15 129.69 127.91 127.02 126.02 124.63 115.49 54.80 54.50 51.60 42.67 41.81 40.46 37.45 36.35 24.59 22.03 20.56 20.08
Parameter Value
1 Title KKS-1-280.100002.fid
2 Origin Bruker BioSpin GmbH
3 Solvent MeOD
4 Temperature 298.2
5 Pulse Sequence zgpg30
6 Experiment 1D
7 Probe Z150363_0002 (CP BBO 500S2 BBF-H-D-05 Z)
8 Number of Scans 5120
9 Receiver Gain 20.2
10 Relaxation Delay 0.1500
11 Pulse Width 9.1000
12 Acquisition Time 1.1010
13 Acquisition Date 2018-10-10T14:33:58
14 Modification Date 2018-10-10T14:33:58
15 Spectrometer Frequency 125.79
16 Spectral Width 29761.9
17 Lowest Frequency -2304.1
18 Nucleus 13C
19 Acquired Size 32768
20 Spectral Size 65536
Me
$^{+}H_3N$
.OAc$^{-}$
OH
1e
Me
Me
f1 (ppm)

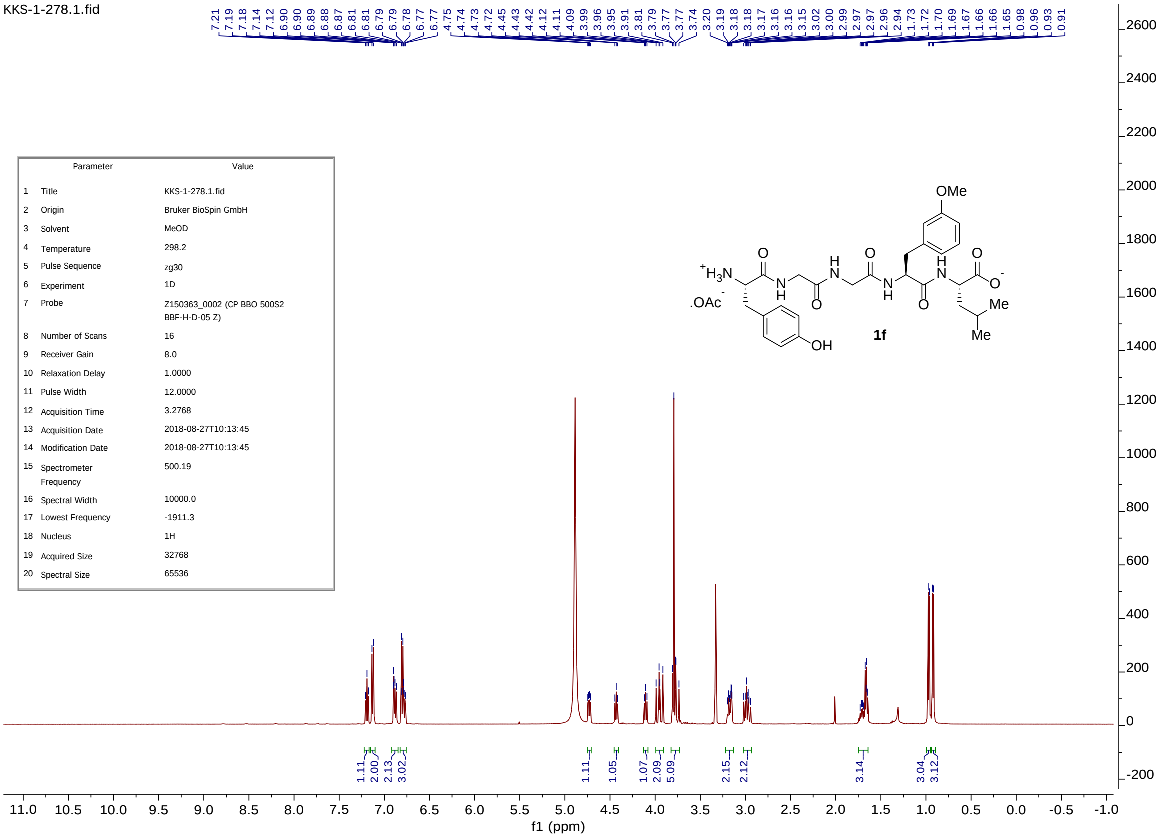


| | Parameter | Value |
|---|---|---|
| 1 | Title | KKS-1-278.1.fid |
| 2 | Origin | Bruker BioSpin GmbH |
| 3 | Solvent | MeOD |
| 4 | Temperature | 298.2 |
| 5 | Pulse Sequence | zg30 |
| 6 | Experiment | 1D |
| 7 | Probe | Z150363_0002 (CP BBO 500S2 BBF-H-D-05 Z) |
| 8 | Number of Scans | 16 |
| 9 | Receiver Gain | 8.0 |
| 10 | Relaxation Delay | 1.0000 |
| 11 | Pulse Width | 12.0000 |
| 12 | Acquisition Time | 3.2768 |
| 13 | Acquisition Date | 2018-08-27T10:13:45 |
| 14 | Modification Date | 2018-08-27T10:13:45 |
| 15 | Spectrometer Frequency | 500.19 |
| 16 | Spectral Width | 10000.0 |
| 17 | Lowest Frequency | -1911.3 |
| 18 | Nucleus | 1H |
| 19 | Acquired Size | 32768 |
| 20 | Spectral Size | 65536 |

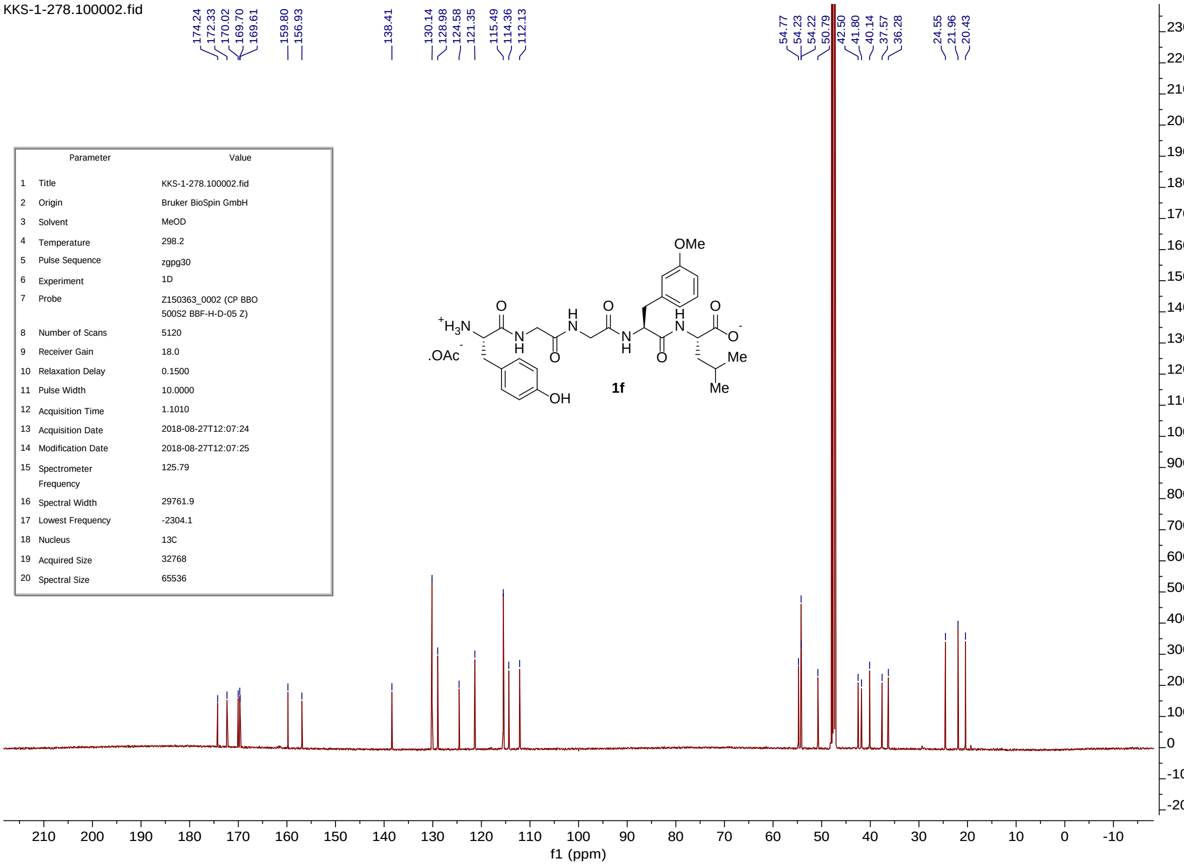


| | Parameter | Value |
|---|---|---|
| 1 | Title | KKS-1-278.100002.fid |
| 2 | Origin | Bruker BioSpin GmbH |
| 3 | Solvent | MeOD |
| 4 | Temperature | 298.2 |
| 5 | Pulse Sequence | zgpg30 |
| 6 | Experiment | 1D |
| 7 | Probe | Z150363_0002 (CP BBO 500S2 BBF-H-D-05 Z) |
| 8 | Number of Scans | 5120 |
| 9 | Receiver Gain | 18.0 |
| 10 | Relaxation Delay | 0.1500 |
| 11 | Pulse Width | 10.0000 |
| 12 | Acquisition Time | 1.1010 |
| 13 | Acquisition Date | 2018-08-27T12:07:24 |
| 14 | Modification Date | 2018-08-27T12:07:25 |
| 15 | Spectrometer Frequency | 125.79 |
| 16 | Spectral Width | 29761.9 |
| 17 | Lowest Frequency | -2304.1 |
| 18 | Nucleus | 13C |
| 19 | Acquired Size | 32768 |
| 20 | Spectral Size | 65536 |

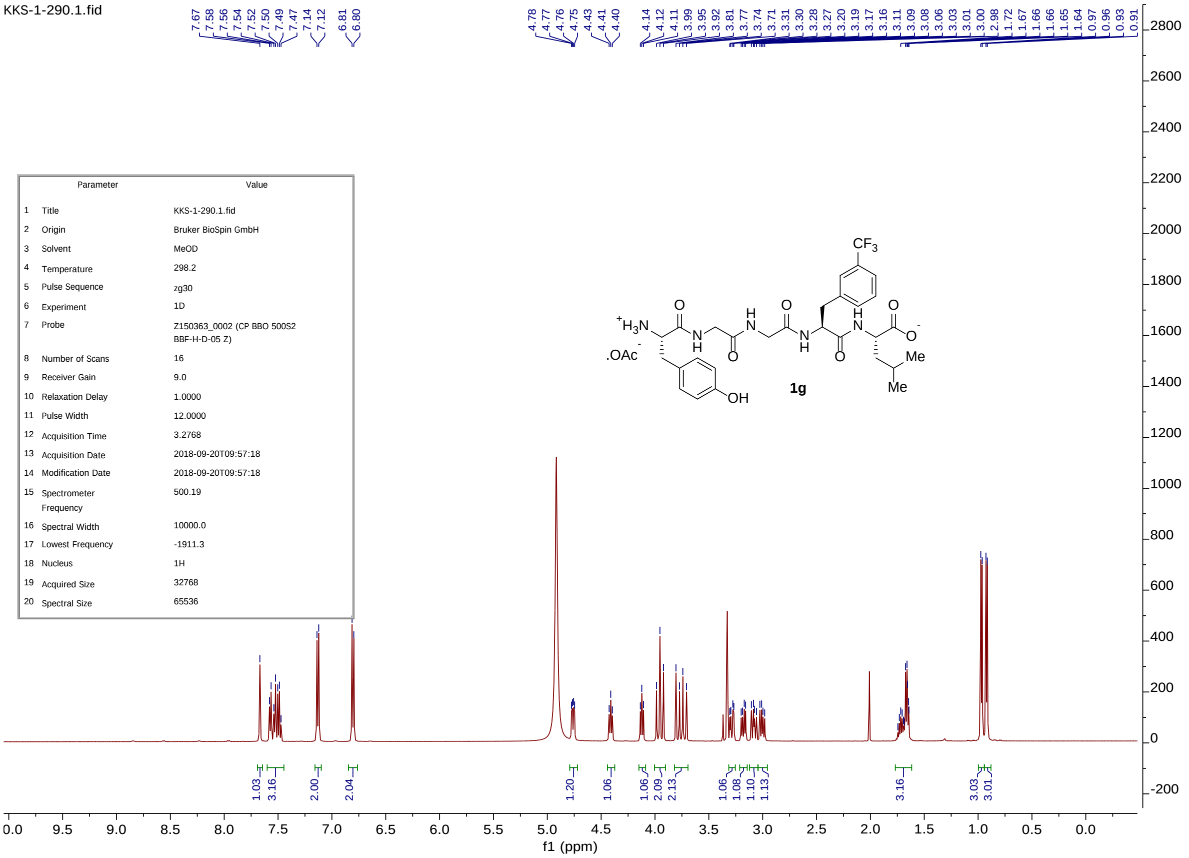


| | Parameter | Value |
|---|---|---|
| 1 | Title | KKS-1-290.1.fid |
| 2 | Origin | Bruker BioSpin GmbH |
| 3 | Solvent | MeOD |
| 4 | Temperature | 298.2 |
| 5 | Pulse Sequence | zg30 |
| 6 | Experiment | 1D |
| 7 | Probe | Z150363_0002 (CP BBO 500S2 BBF-H-D-05 Z) |
| 8 | Number of Scans | 16 |
| 9 | Receiver Gain | 9.0 |
| 10 | Relaxation Delay | 1.0000 |
| 11 | Pulse Width | 12.0000 |
| 12 | Acquisition Time | 3.2768 |
| 13 | Acquisition Date | 2018-09-20T09:57:18 |
| 14 | Modification Date | 2018-09-20T09:57:18 |
| 15 | Spectrometer Frequency | 500.19 |
| 16 | Spectral Width | 10000.0 |
| 17 | Lowest Frequency | -1911.3 |
| 18 | Nucleus | 1H |
| 19 | Acquired Size | 32768 |
| 20 | Spectral Size | 65536 |

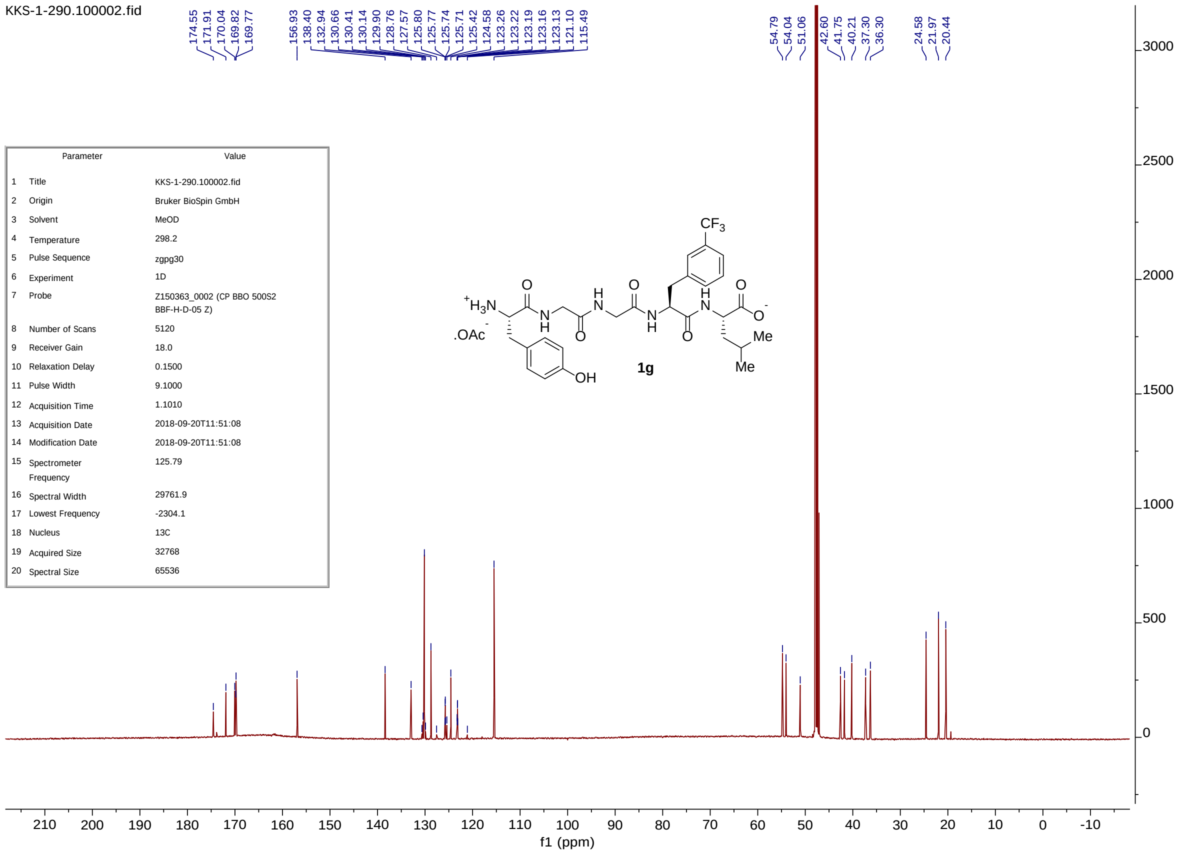


| | Parameter | Value |
|---|---|---|
| 1 | Title | KKS-1-290.100002.fid |
| 2 | Origin | Bruker BioSpin GmbH |
| 3 | Solvent | MeOD |
| 4 | Temperature | 298.2 |
| 5 | Pulse Sequence | zgpg30 |
| 6 | Experiment | 1D |
| 7 | Probe | Z150363_0002 (CP BBO 500S2 BBF-H-D-05 Z) |
| 8 | Number of Scans | 5120 |
| 9 | Receiver Gain | 18.0 |
| 10 | Relaxation Delay | 0.1500 |
| 11 | Pulse Width | 9.1000 |
| 12 | Acquisition Time | 1.1010 |
| 13 | Acquisition Date | 2018-09-20T11:51:08 |
| 14 | Modification Date | 2018-09-20T11:51:08 |
| 15 | Spectrometer Frequency | 125.79 |
| 16 | Spectral Width | 29761.9 |
| 17 | Lowest Frequency | -2304.1 |
| 18 | Nucleus | 13C |
| 19 | Acquired Size | 32768 |
| 20 | Spectral Size | 65536 |

kks-1-290_fr4.1.fid

| | Parameter | Value |
|---|---|---|
| 1 | Title | kks-1-290_fr4.1.fid |
| 2 | Origin | Bruker Analytik GmbH |
| 3 | Solvent | None |
| 4 | Temperature | 297.2 |
| 5 | Pulse Sequence | zgflqn |
| 6 | Experiment | 1D |
| 7 | Probe | 5 mm PABBO BB/ 19F-1H/ D Z-GRD Z110902/ 0011 |
| 8 | Number of Scans | 16 |
| 9 | Receiver Gain | 812.7 |
| 10 | Relaxation Delay | 1.0000 |
| 11 | Pulse Width | 19.7000 |
| 12 | Acquisition Time | 0.6554 |
| 13 | Acquisition Date | 2019-07-12T18:24:18 |
| 14 | Modification Date | 2019-07-12T18:24:21 |
| 15 | Spectrometer Frequency | 470.50 |
| 16 | Spectral Width | 100000.0 |
| 17 | Lowest Frequency | -96974.3 |
| 18 | Nucleus | 19F |
| 19 | Acquired Size | 65536 |
| 20 | Spectral Size | 131072 |

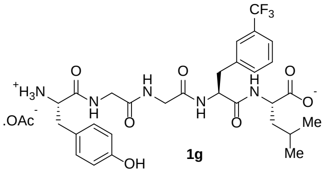

**1g**

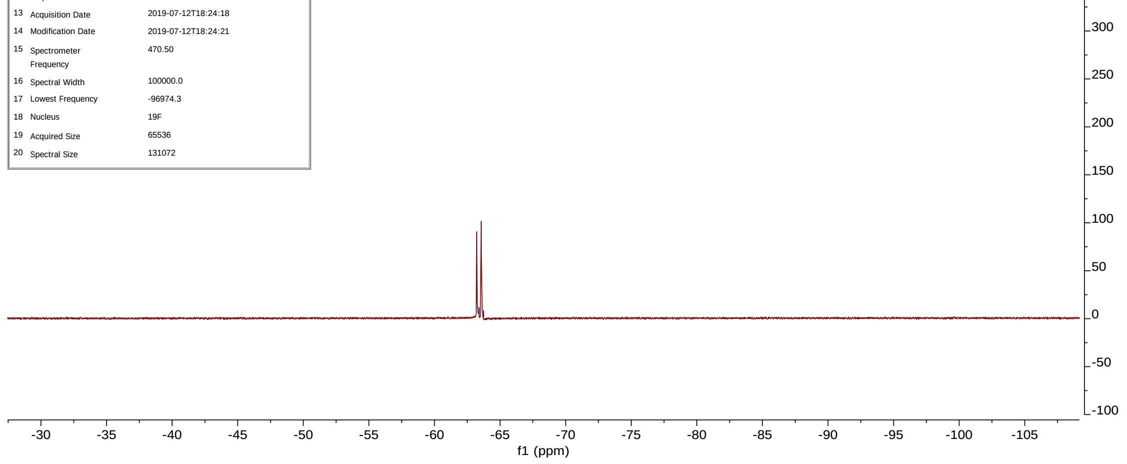

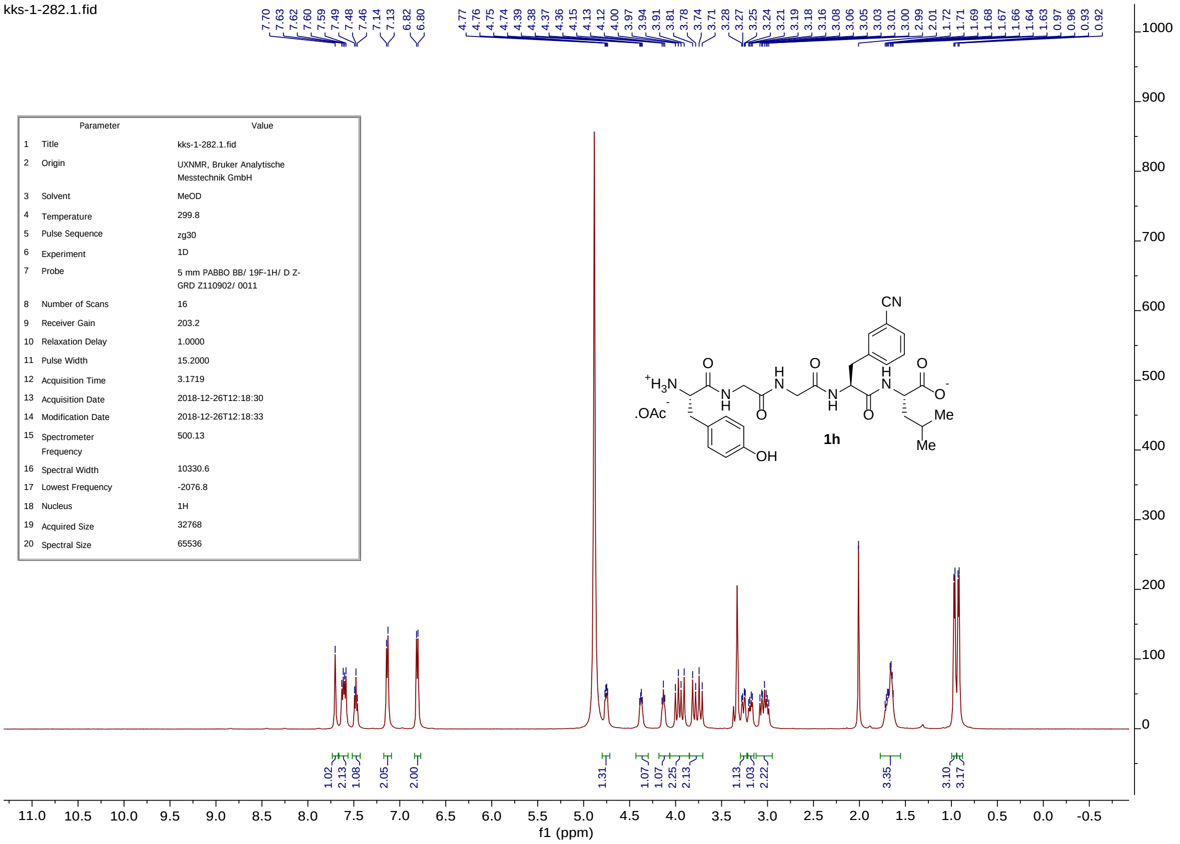
kks-1-282.1.fid
7.70 7.63 7.62 7.60 7.59 7.49 7.48 7.46 7.14 7.13 6.82 6.80 4.77 4.76 4.75 4.74 4.39 4.38 4.37 4.36 4.15 4.13 4.12 4.00 3.97 3.94 3.91 3.81 3.78 3.74 3.71 3.28 3.27 3.25 3.24 3.21 3.19 3.18 3.16 3.08 3.06 3.05 3.03 3.01 3.00 2.99 2.01 1.72 1.71 1.69 1.68 1.67 1.66 1.64 1.63 0.97 0.96 0.93 0.92
Parameter Value
1 Title kks-1-282.1.fid
2 Origin UXNMR, Bruker Analytische Messtechnik GmbH
3 Solvent MeOD
4 Temperature 299.8
5 Pulse Sequence zg30
6 Experiment 1D
7 Probe 5 mm PABBO BB/ 19F-1H/ D Z-GRD Z110902/ 0011
8 Number of Scans 16
9 Receiver Gain 203.2
10 Relaxation Delay 1.0000
11 Pulse Width 15.2000
12 Acquisition Time 3.1719
13 Acquisition Date 2018-12-26T12:18:30
14 Modification Date 2018-12-26T12:18:33
15 Spectrometer Frequency 500.13
16 Spectral Width 10330.6
17 Lowest Frequency -2076.8
18 Nucleus 1H
19 Acquired Size 32768
20 Spectral Size 65536
CN
+H3N
.OAc
OH
Me
Me
1h
1.02 2.13 1.08 2.05 2.00 1.31 1.07 1.07 2.25 2.13 1.13 1.03 2.22 3.35 3.10 3.17
f1 (ppm)

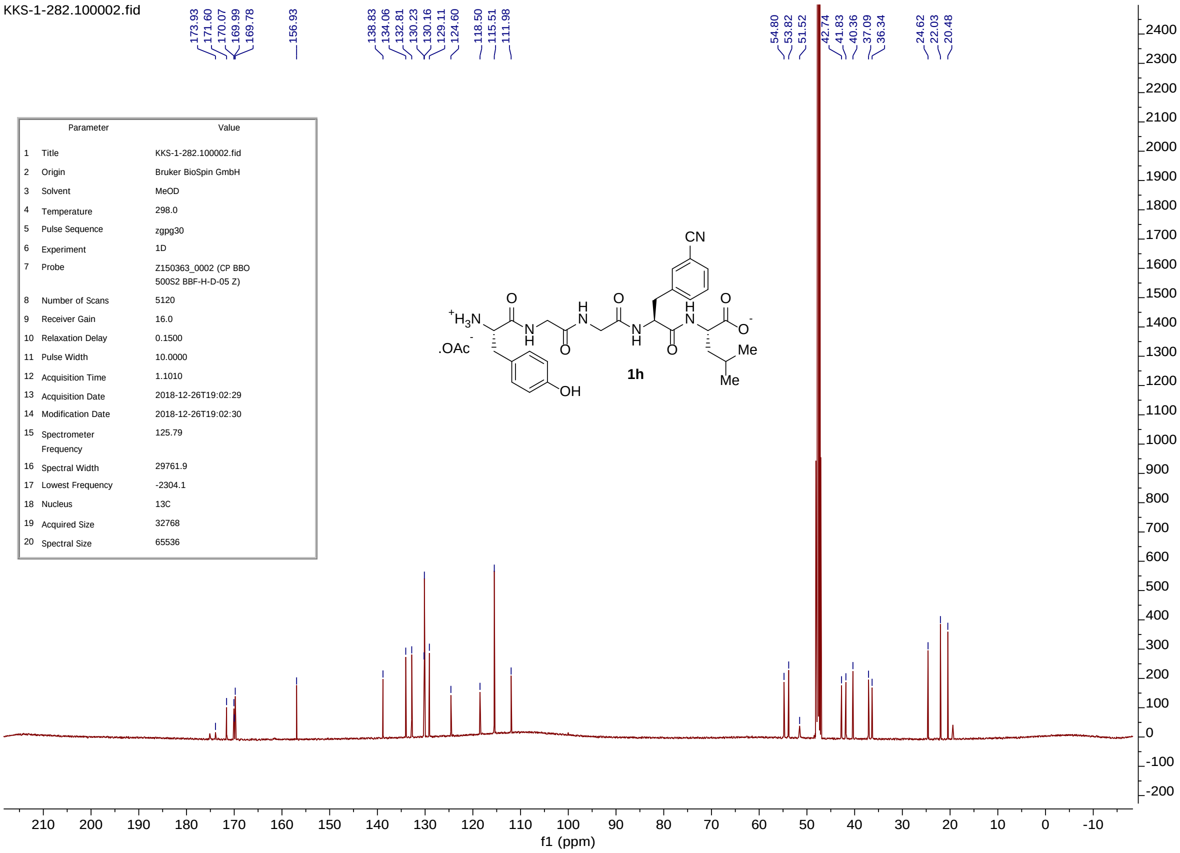
KKS-1-282.100002.fid
173.93 171.60 170.07 169.99 169.78 156.93 138.83 134.06 132.81 130.23 130.16 129.11 124.60 118.50 115.51 111.98 54.80 53.82 51.52 42.74 41.83 40.36 37.09 36.34 24.62 22.03 20.48
Parameter Value
1 Title KKS-1-282.100002.fid
2 Origin Bruker BioSpin GmbH
3 Solvent MeOD
4 Temperature 298.0
5 Pulse Sequence zgpg30
6 Experiment 1D
7 Probe Z150363_0002 (CP BBO 500S2 BBF-H-D-05 Z)
8 Number of Scans 5120
9 Receiver Gain 16.0
10 Relaxation Delay 0.1500
11 Pulse Width 10.0000
12 Acquisition Time 1.1010
13 Acquisition Date 2018-12-26T19:02:29
14 Modification Date 2018-12-26T19:02:30
15 Spectrometer Frequency 125.79
16 Spectral Width 29761.9
17 Lowest Frequency -2304.1
18 Nucleus 13C
19 Acquired Size 32768
20 Spectral Size 65536
CN
+H3N
.OAc
OH
Me
Me
1h
f1 (ppm)

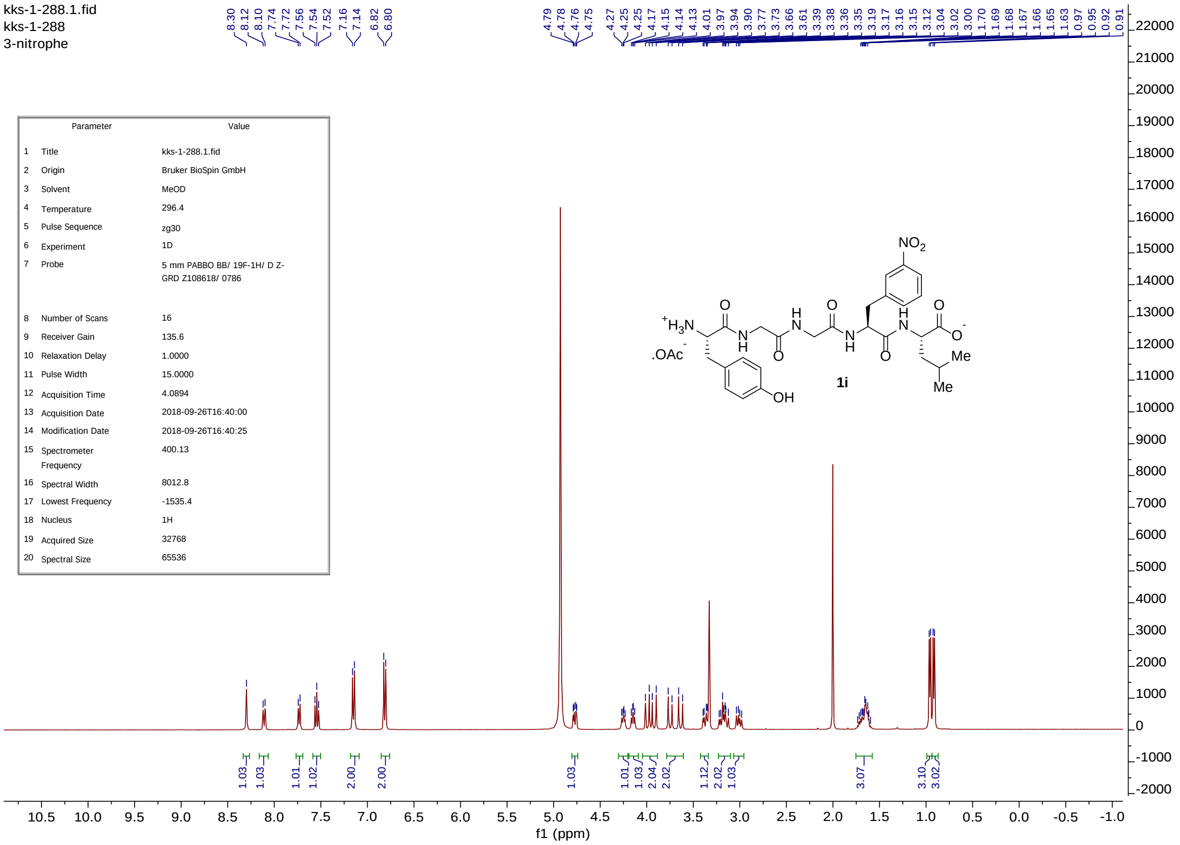


| | Parameter | Value |
|---|---|---|
| 1 | Title | kks-1-288.1.fid |
| 2 | Origin | Bruker BioSpin GmbH |
| 3 | Solvent | MeOD |
| 4 | Temperature | 296.4 |
| 5 | Pulse Sequence | zg30 |
| 6 | Experiment | 1D |
| 7 | Probe | 5 mm PABBO BB/ 19F-1H/ D Z-GRD Z108618/ 0786 |
| 8 | Number of Scans | 16 |
| 9 | Receiver Gain | 135.6 |
| 10 | Relaxation Delay | 1.0000 |
| 11 | Pulse Width | 15.0000 |
| 12 | Acquisition Time | 4.0894 |
| 13 | Acquisition Date | 2018-09-26T16:40:00 |
| 14 | Modification Date | 2018-09-26T16:40:25 |
| 15 | Spectrometer Frequency | 400.13 |
| 16 | Spectral Width | 8012.8 |
| 17 | Lowest Frequency | -1535.4 |
| 18 | Nucleus | 1H |
| 19 | Acquired Size | 32768 |
| 20 | Spectral Size | 65536 |

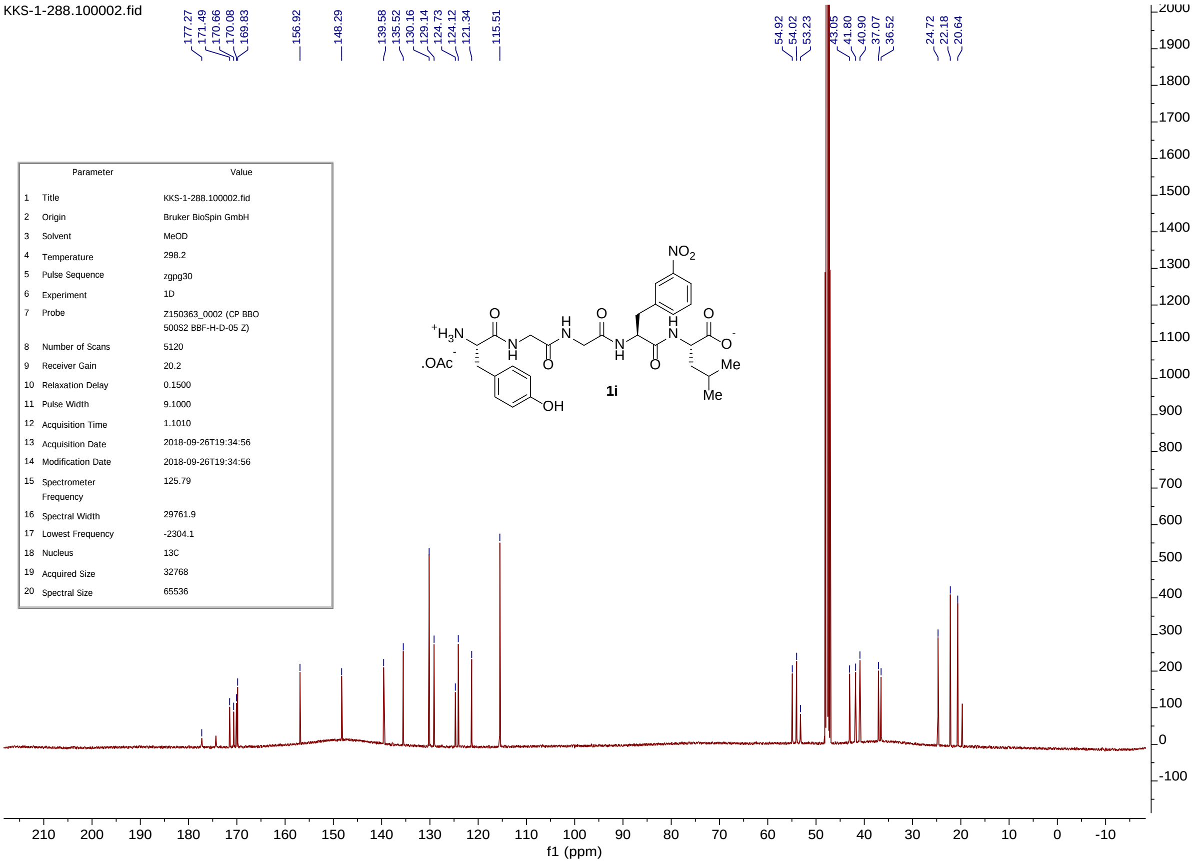


| | Parameter | Value |
|---|---|---|
| 1 | Title | KKS-1-288.100002.fid |
| 2 | Origin | Bruker BioSpin GmbH |
| 3 | Solvent | MeOD |
| 4 | Temperature | 298.2 |
| 5 | Pulse Sequence | zgpg30 |
| 6 | Experiment | 1D |
| 7 | Probe | Z150363_0002 (CP BBO 500S2 BBF-H-D-05 Z) |
| 8 | Number of Scans | 5120 |
| 9 | Receiver Gain | 20.2 |
| 10 | Relaxation Delay | 0.1500 |
| 11 | Pulse Width | 9.1000 |
| 12 | Acquisition Time | 1.1010 |
| 13 | Acquisition Date | 2018-09-26T19:34:56 |
| 14 | Modification Date | 2018-09-26T19:34:56 |
| 15 | Spectrometer Frequency | 125.79 |
| 16 | Spectral Width | 29761.9 |
| 17 | Lowest Frequency | -2304.1 |
| 18 | Nucleus | 13C |
| 19 | Acquired Size | 32768 |
| 20 | Spectral Size | 65536 |

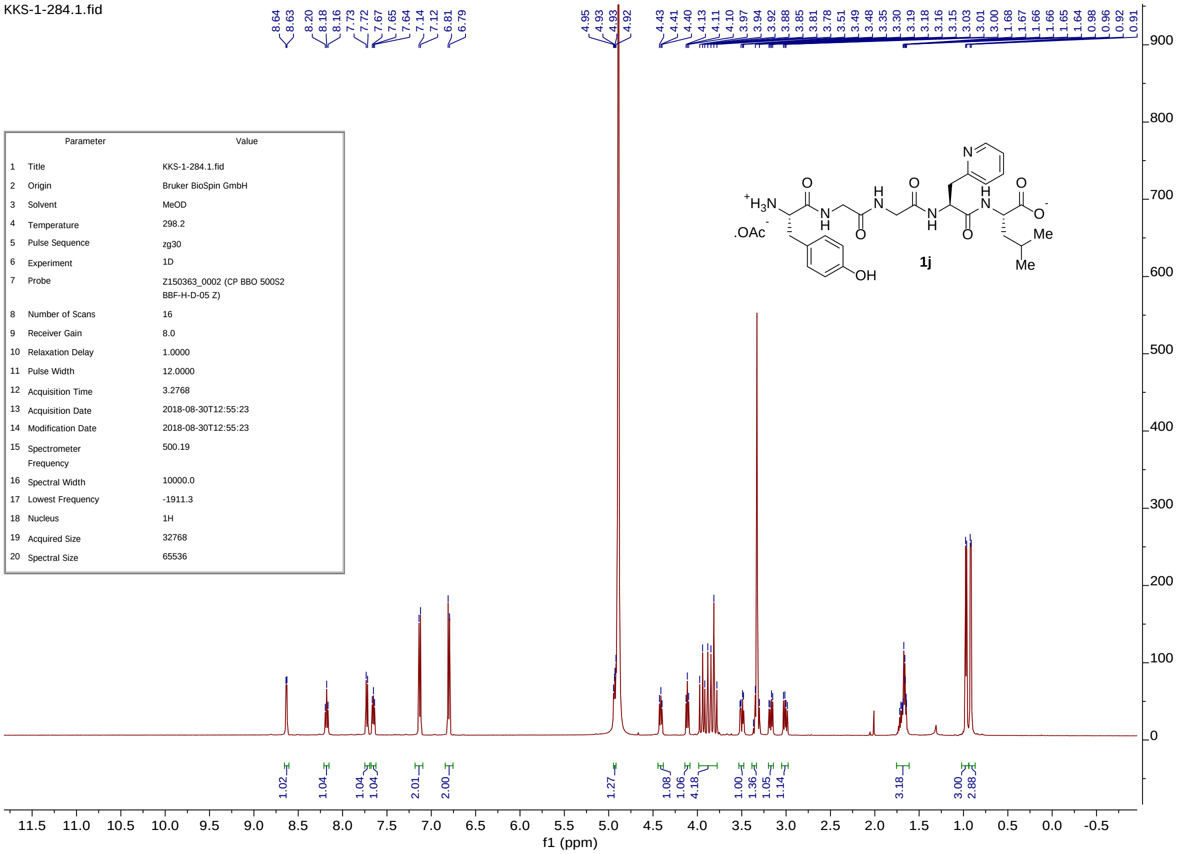


| | Parameter | Value |
|---|---|---|
| 1 | Title | KKS-1-284.1.fid |
| 2 | Origin | Bruker BioSpin GmbH |
| 3 | Solvent | MeOD |
| 4 | Temperature | 298.2 |
| 5 | Pulse Sequence | zg30 |
| 6 | Experiment | 1D |
| 7 | Probe | Z150363_0002 (CP BBO 500S2 BBF-H-D-05 Z) |
| 8 | Number of Scans | 16 |
| 9 | Receiver Gain | 8.0 |
| 10 | Relaxation Delay | 1.0000 |
| 11 | Pulse Width | 12.0000 |
| 12 | Acquisition Time | 3.2768 |
| 13 | Acquisition Date | 2018-08-30T12:55:23 |
| 14 | Modification Date | 2018-08-30T12:55:23 |
| 15 | Spectrometer Frequency | 500.19 |
| 16 | Spectral Width | 10000.0 |
| 17 | Lowest Frequency | -1911.3 |
| 18 | Nucleus | 1H |
| 19 | Acquired Size | 32768 |
| 20 | Spectral Size | 65536 |

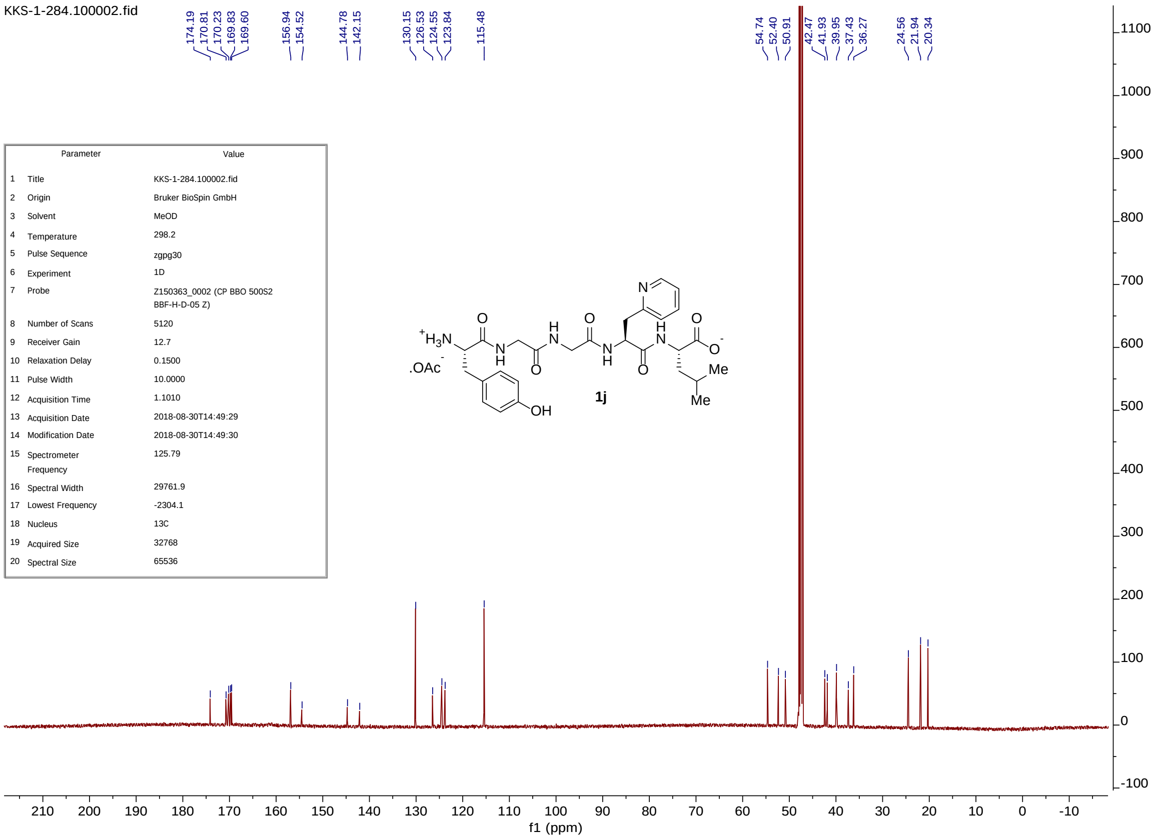


| | Parameter | Value |
|---|---|---|
| 1 | Title | KKS-1-284.100002.fid |
| 2 | Origin | Bruker BioSpin GmbH |
| 3 | Solvent | MeOD |
| 4 | Temperature | 298.2 |
| 5 | Pulse Sequence | zgpg30 |
| 6 | Experiment | 1D |
| 7 | Probe | Z150363_0002 (CP BBO 500S2 BBF-H-D-05 Z) |
| 8 | Number of Scans | 5120 |
| 9 | Receiver Gain | 12.7 |
| 10 | Relaxation Delay | 0.1500 |
| 11 | Pulse Width | 10.0000 |
| 12 | Acquisition Time | 1.1010 |
| 13 | Acquisition Date | 2018-08-30T14:49:29 |
| 14 | Modification Date | 2018-08-30T14:49:30 |
| 15 | Spectrometer Frequency | 125.79 |
| 16 | Spectral Width | 29761.9 |
| 17 | Lowest Frequency | -2304.1 |
| 18 | Nucleus | 13C |
| 19 | Acquired Size | 32768 |
| 20 | Spectral Size | 65536 |

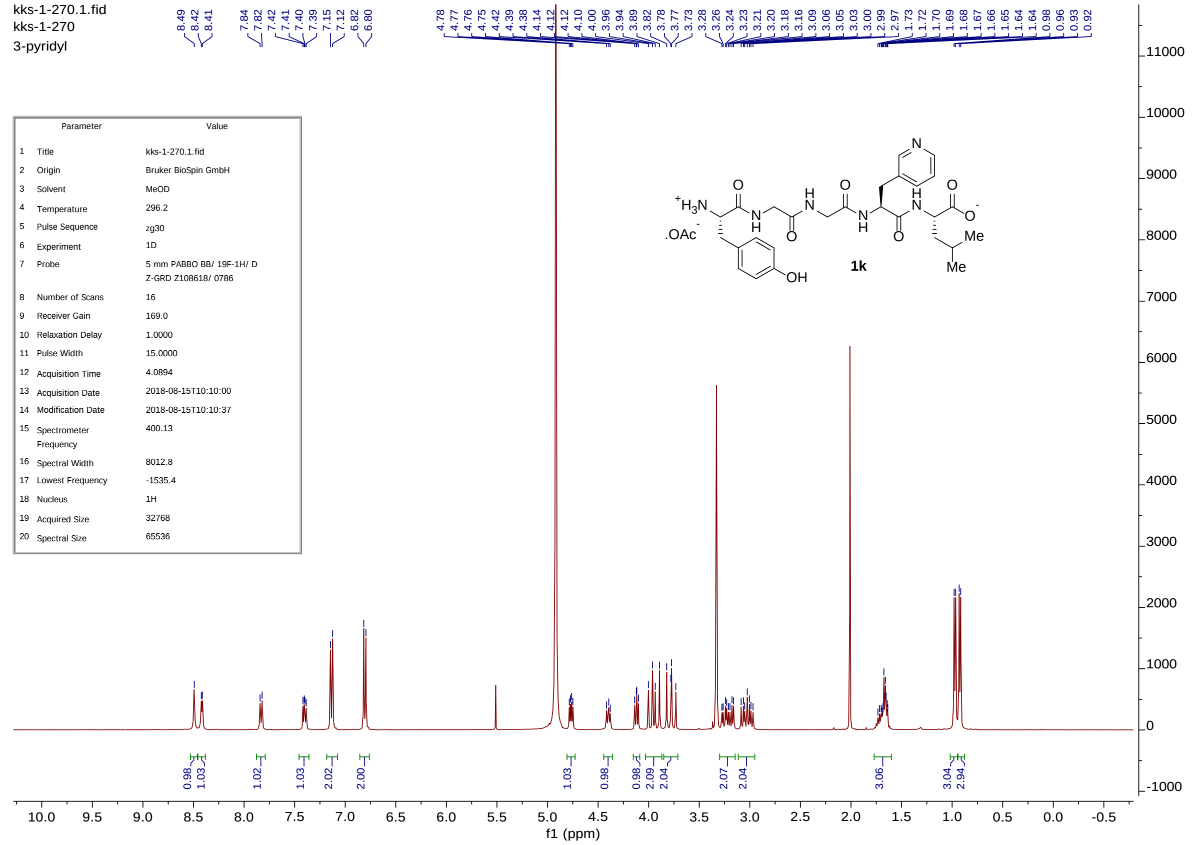


| | Parameter | Value |
|---|---|---|
| 1 | Title | kks-1-270.1.fid |
| 2 | Origin | Bruker BioSpin GmbH |
| 3 | Solvent | MeOD |
| 4 | Temperature | 296.2 |
| 5 | Pulse Sequence | zg30 |
| 6 | Experiment | 1D |
| 7 | Probe | 5 mm PABBO BB/ 19F-1H/ D Z-GRD Z108618/ 0786 |
| 8 | Number of Scans | 16 |
| 9 | Receiver Gain | 169.0 |
| 10 | Relaxation Delay | 1.0000 |
| 11 | Pulse Width | 15.0000 |
| 12 | Acquisition Time | 4.0894 |
| 13 | Acquisition Date | 2018-08-15T10:10:00 |
| 14 | Modification Date | 2018-08-15T10:10:37 |
| 15 | Spectrometer Frequency | 400.13 |
| 16 | Spectral Width | 8012.8 |
| 17 | Lowest Frequency | -1535.4 |
| 18 | Nucleus | 1H |
| 19 | Acquired Size | 32768 |
| 20 | Spectral Size | 65536 |

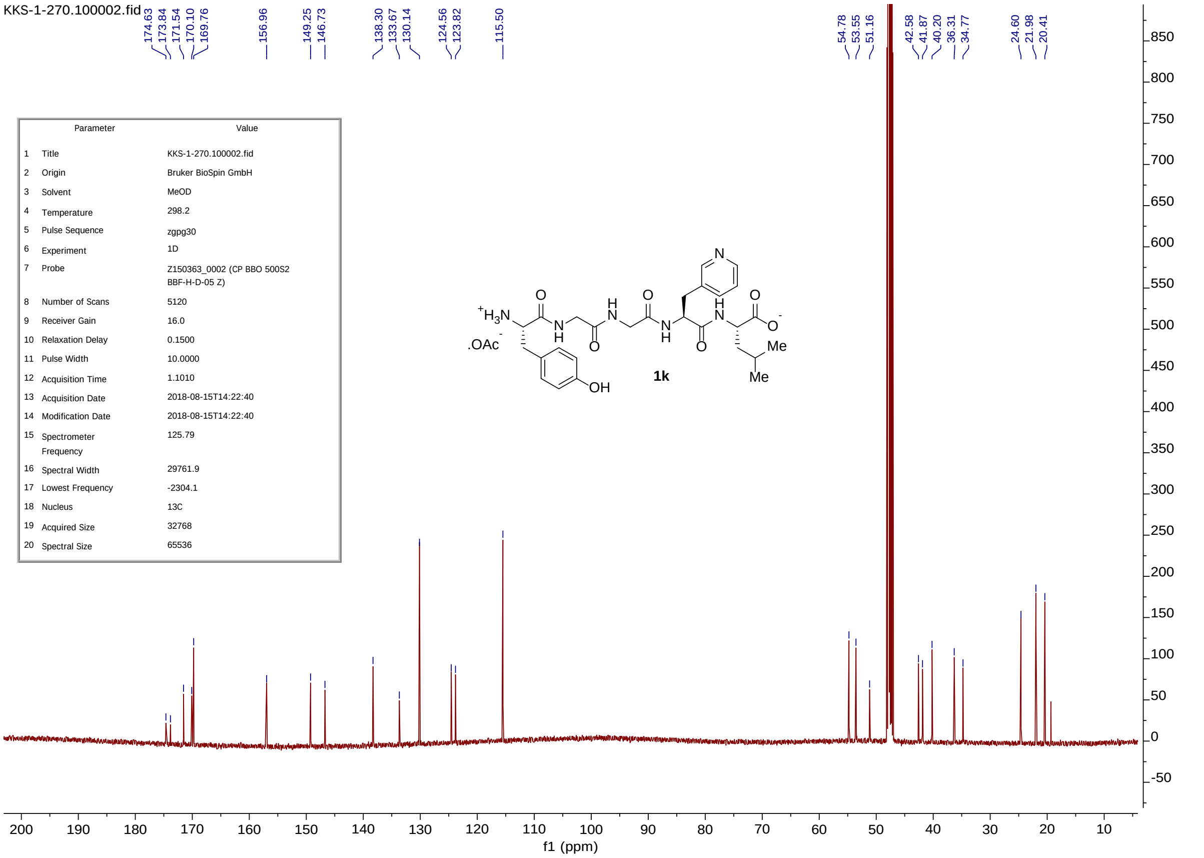


| | Parameter | Value |
|---|---|---|
| 1 | Title | KKS-1-270.100002.fid |
| 2 | Origin | Bruker BioSpin GmbH |
| 3 | Solvent | MeOD |
| 4 | Temperature | 298.2 |
| 5 | Pulse Sequence | zgpg30 |
| 6 | Experiment | 1D |
| 7 | Probe | Z150363_0002 (CP BBO 500S2 BBF-H-D-05 Z) |
| 8 | Number of Scans | 5120 |
| 9 | Receiver Gain | 16.0 |
| 10 | Relaxation Delay | 0.1500 |
| 11 | Pulse Width | 10.0000 |
| 12 | Acquisition Time | 1.1010 |
| 13 | Acquisition Date | 2018-08-15T14:22:40 |
| 14 | Modification Date | 2018-08-15T14:22:40 |
| 15 | Spectrometer Frequency | 125.79 |
| 16 | Spectral Width | 29761.9 |
| 17 | Lowest Frequency | -2304.1 |
| 18 | Nucleus | 13C |
| 19 | Acquired Size | 32768 |
| 20 | Spectral Size | 65536 |

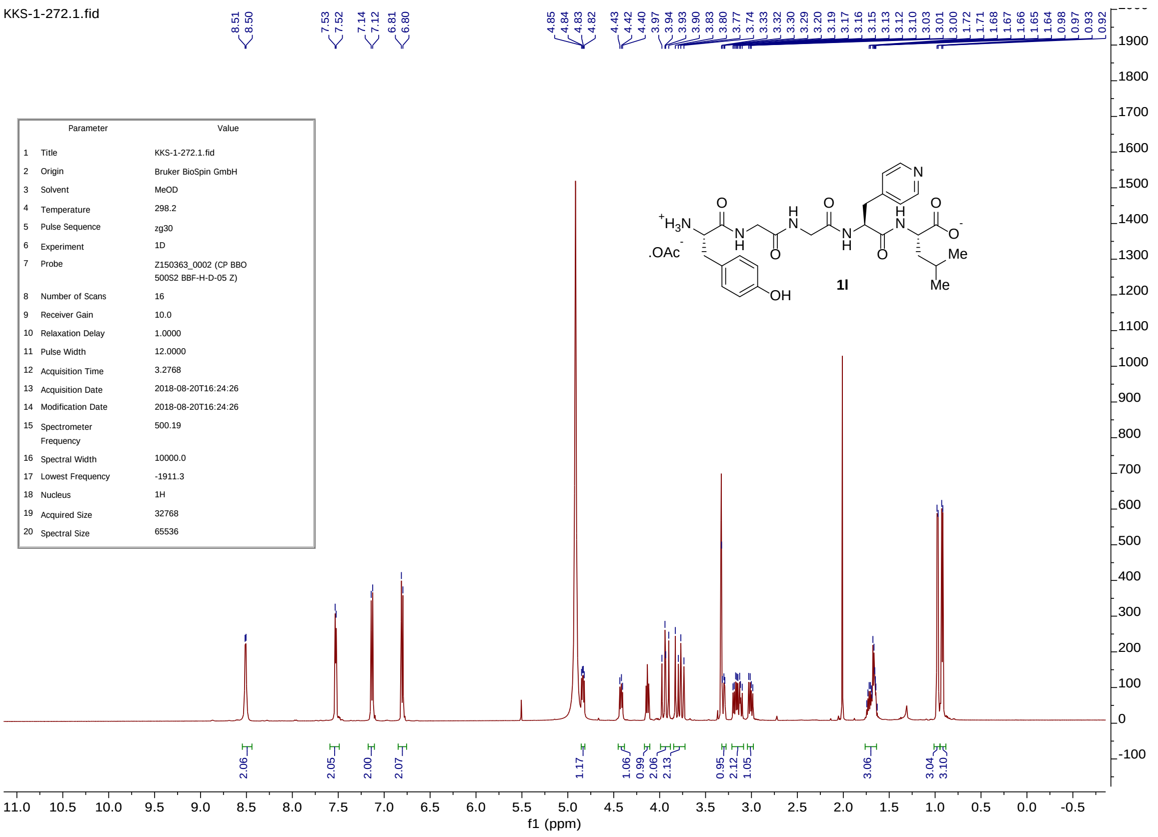


| | Parameter | Value |
|---|---|---|
| 1 | Title | KKS-1-272.1.fid |
| 2 | Origin | Bruker BioSpin GmbH |
| 3 | Solvent | MeOD |
| 4 | Temperature | 298.2 |
| 5 | Pulse Sequence | zg30 |
| 6 | Experiment | 1D |
| 7 | Probe | Z150363_0002 (CP BBO 500S2 BBF-H-D-05 Z) |
| 8 | Number of Scans | 16 |
| 9 | Receiver Gain | 10.0 |
| 10 | Relaxation Delay | 1.0000 |
| 11 | Pulse Width | 12.0000 |
| 12 | Acquisition Time | 3.2768 |
| 13 | Acquisition Date | 2018-08-20T16:24:26 |
| 14 | Modification Date | 2018-08-20T16:24:26 |
| 15 | Spectrometer Frequency | 500.19 |
| 16 | Spectral Width | 10000.0 |
| 17 | Lowest Frequency | -1911.3 |
| 18 | Nucleus | 1H |
| 19 | Acquired Size | 32768 |
| 20 | Spectral Size | 65536 |

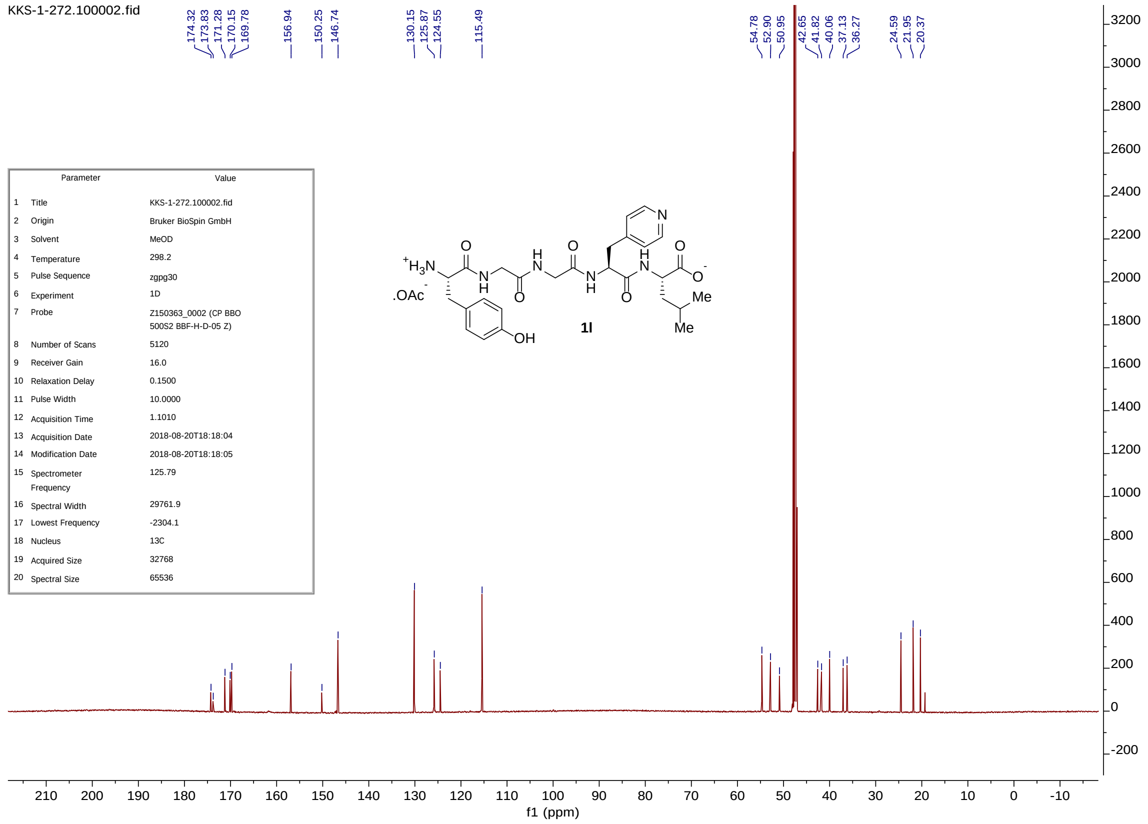


| | Parameter | Value |
|---|---|---|
| 1 | Title | KKS-1-272.100002.fid |
| 2 | Origin | Bruker BioSpin GmbH |
| 3 | Solvent | MeOD |
| 4 | Temperature | 298.2 |
| 5 | Pulse Sequence | zgpg30 |
| 6 | Experiment | 1D |
| 7 | Probe | Z150363_0002 (CP BBO 500S2 BBF-H-D-05 Z) |
| 8 | Number of Scans | 5120 |
| 9 | Receiver Gain | 16.0 |
| 10 | Relaxation Delay | 0.1500 |
| 11 | Pulse Width | 10.0000 |
| 12 | Acquisition Time | 1.1010 |
| 13 | Acquisition Date | 2018-08-20T18:18:04 |
| 14 | Modification Date | 2018-08-20T18:18:05 |
| 15 | Spectrometer Frequency | 125.79 |
| 16 | Spectral Width | 29761.9 |
| 17 | Lowest Frequency | -2304.1 |
| 18 | Nucleus | 13C |
| 19 | Acquired Size | 32768 |
| 20 | Spectral Size | 65536 |

#### HPLC Chromatograms of Peptides

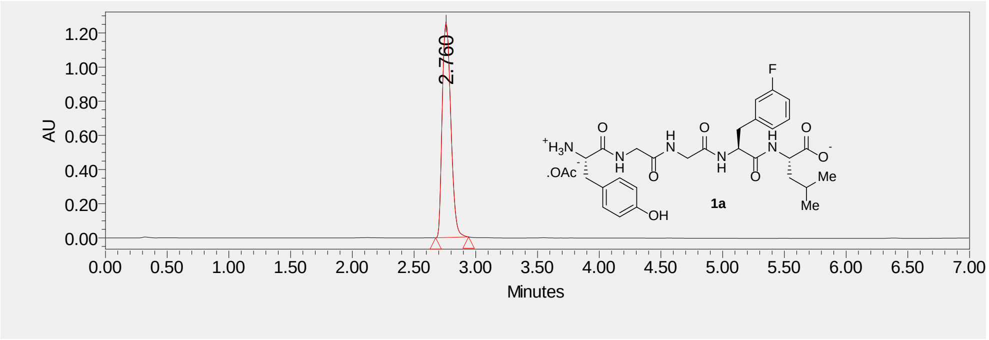


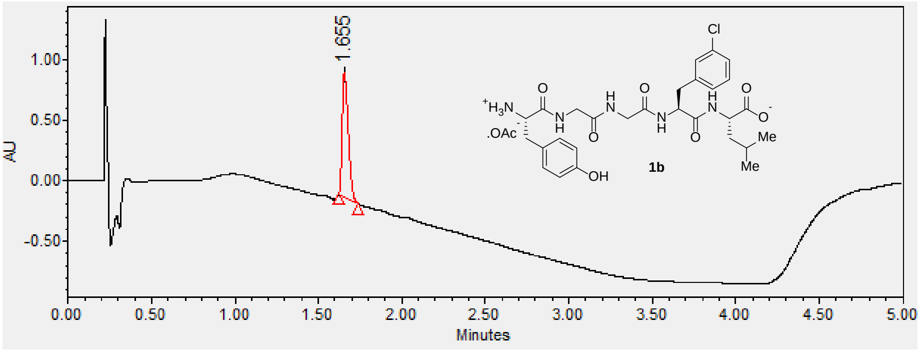


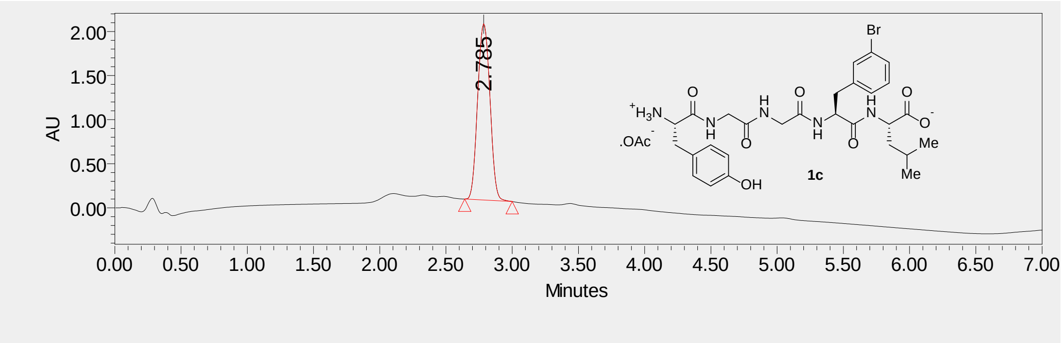

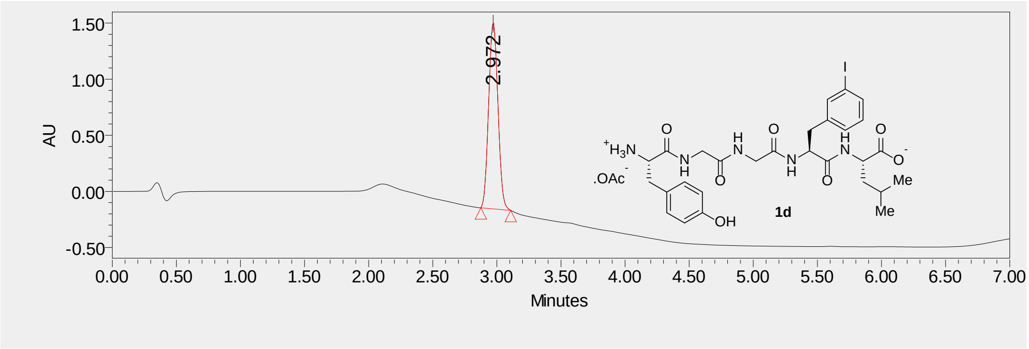
1.50
1.00
0.50
0.00
-0.50
AU
2.972
0.00 0.50 1.00 1.50 2.00 2.50 3.00 3.50 4.00 4.50 5.00 5.50 6.00 6.50 7.00
Minutes
1d

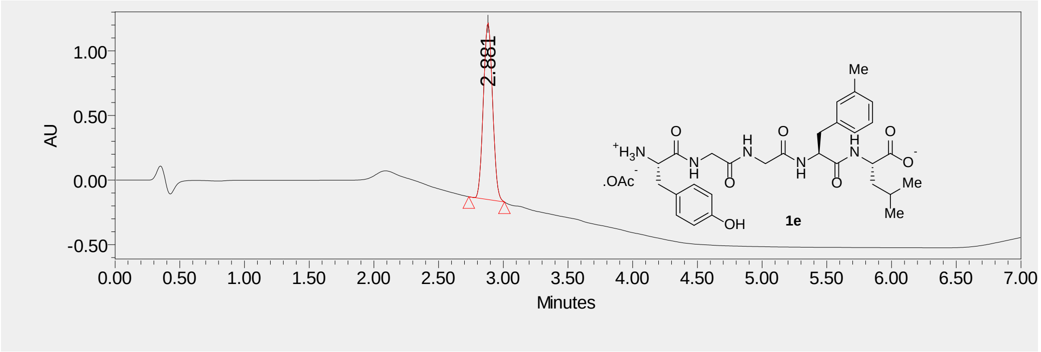
1.00
0.50
0.00
-0.50
AU
2.881
0.00 0.50 1.00 1.50 2.00 2.50 3.00 3.50 4.00 4.50 5.00 5.50 6.00 6.50 7.00
Minutes
1e

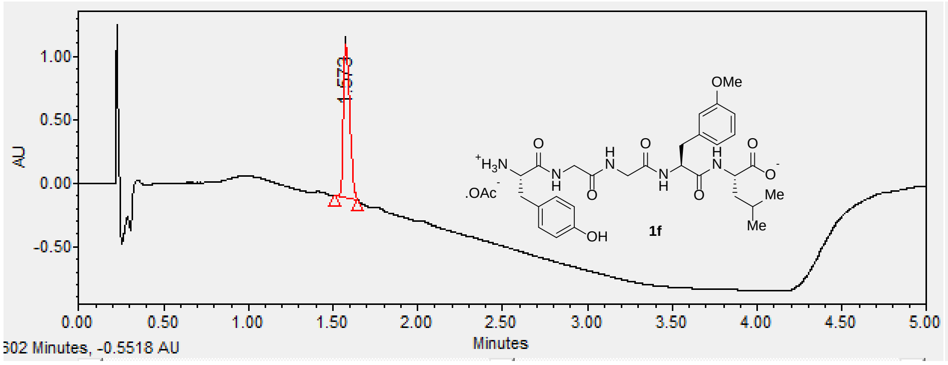
1.00
0.50
0.00
-0.50
AU
1.573
0.00 0.50 1.00 1.50 2.00 2.50 3.00 3.50 4.00 4.50 5.00
Minutes
502 Minutes, -0.5518 AU
1f

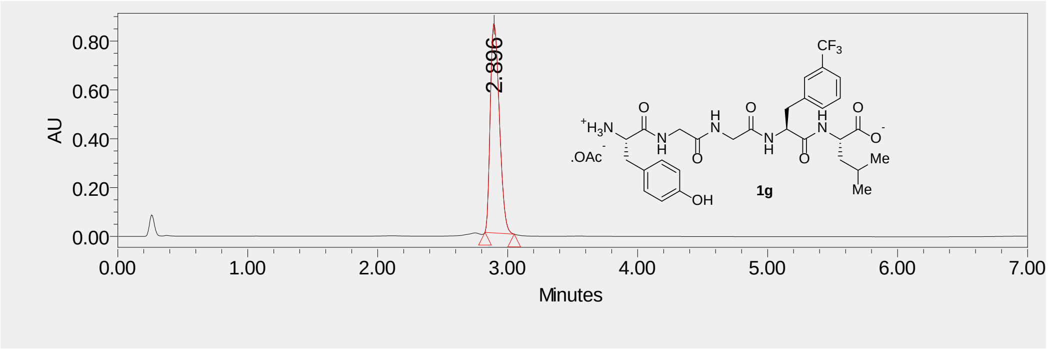
0.80
0.60
0.40
0.20
0.00
AU
2.896
0.00
1.00
2.00
3.00
4.00
5.00
6.00
7.00
Minutes
CF3
+H3N
.OAc-
O
N
H
OH
1g
Me
Me

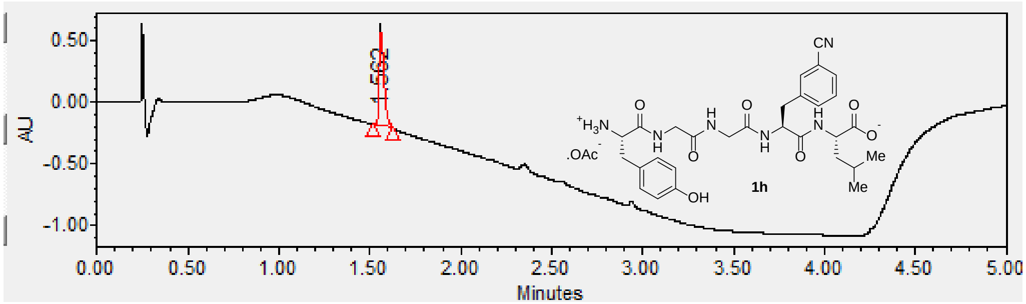
0.50
0.00
-0.50
-1.00
AU
0.00
0.50
1.00
1.50
2.00
2.50
3.00
3.50
4.00
4.50
5.00
Minutes
CN
+H3N
.OAc-
OH
1h
Me
Me

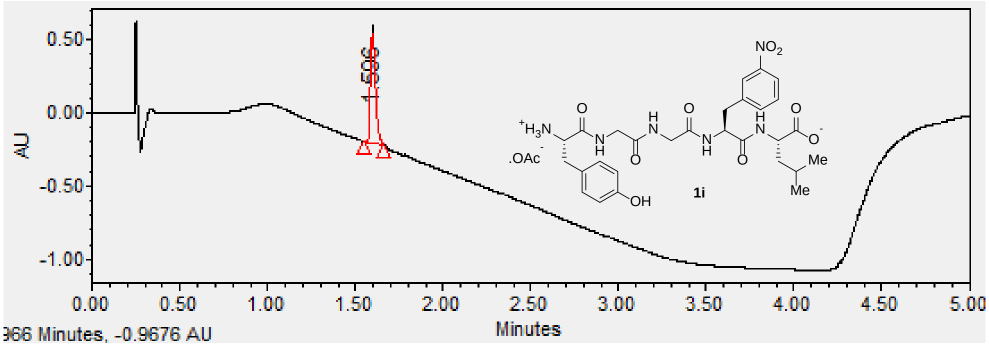
0.50
0.00
-0.50
-1.00
AU
0.00
0.50
1.00
1.50
2.00
2.50
3.00
3.50
4.00
4.50
5.00
Minutes
966 Minutes, -0.9676 AU
NO2
+H3N
.OAc-
OH
1i
Me
Me

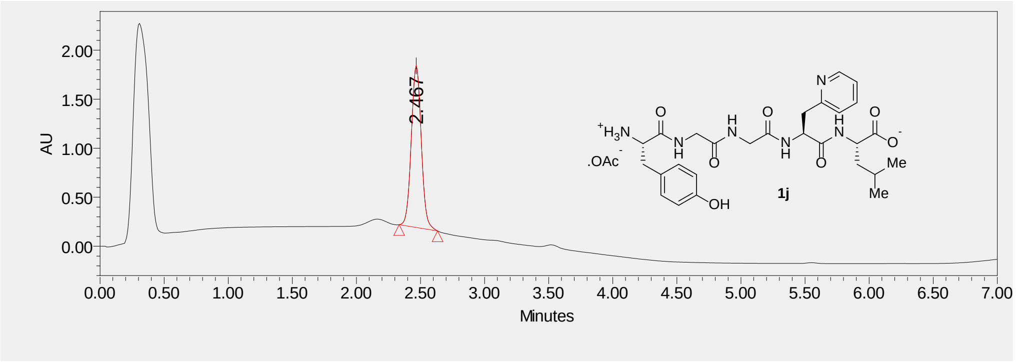
2.00
1.50
1.00
0.50
0.00
AU
2.467
0.00
0.50
1.00
1.50
2.00
2.50
3.00
3.50
4.00
4.50
5.00
5.50
6.00
6.50
7.00
Minutes
1j

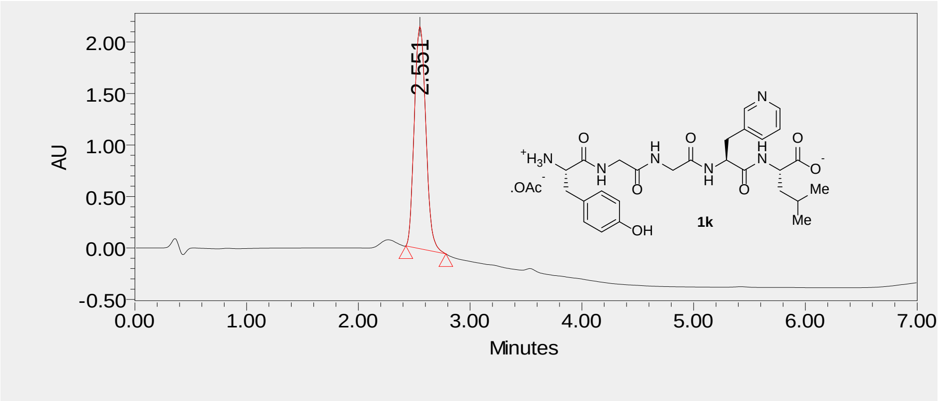
2.00
1.50
1.00
0.50
0.00
-0.50
AU
2.551
0.00
1.00
2.00
3.00
4.00
5.00
6.00
7.00
Minutes
1k

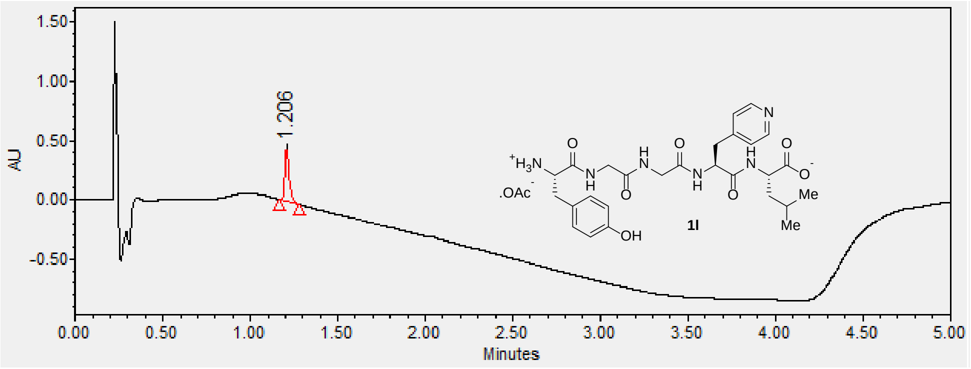
1.50
1.00
0.50
0.00
-0.50
AU
1.206
0.00
0.50
1.00
1.50
2.00
2.50
3.00
3.50
4.00
4.50
5.00
Minutes
1l
